# Expanding the space of self-reproducing ribozymes using probabilistic generative models

**DOI:** 10.1101/2024.07.31.605758

**Authors:** Camille N. Lambert, Vaitea Opuu, Francesco Calvanese, Francesco Zamponi, Eric Hayden, Martin Weigt, Matteo Smerlak, Philippe Nghe

## Abstract

Estimating the plausibility of RNA self-reproduction is central to origin-of-life scenarios but self-reproduction has been shown in only a handful of systems. Here, we populated a vast sequence space of ribozymes using statistical covariation models and secondary structure prediction. Experimentally assayed sequences were found active as far as 65 mutations from a reference natural sequence. The number of potentially generated sequences together with the experimental success rate indicate that at least ∼10^39^ such ribozymes may exist. Randomly sampled artificial ribozymes exhibited autocatalytic self-reproduction akin to the reference sequence. The combination of high-throughput screening and probabilistic modeling considerably improves our estimation of the number of self-reproducing systems, paving the way for a statistical approach to the origin of life.

## Main Text

Origin of life studies have so far focused on finding single instances of molecules involved in abiogenesis. However, assessing the plausibility of the origin of life requires estimating the probability of these molecules to appear. One major scenario is that of the RNA World, which hypothesizes that RNA molecules preceded DNA replication and coded protein synthesis (*1*). Support for this theory has come through the engineering of RNA molecules that catalyze RNA-dependent RNA polymerization (*2, 3*). However, the currently known RNA polymerase ribozymes are too large and complex for self-reproduction. Instead of starting from polymerase ribozymes, self-reproduction may have emerged gradually by the ligation of short RNA oligos to create larger ribozymes promoting their own production (*4*). Such mode of self-reproduction is termed autocatalytic and has been demonstrated in two experimental systems: the R3C mutual ligase (*5*) and the fragmented Azoarcus Group I Intron ribozyme (here denoted Azo) (*6*). Both systems have been diversified into multiple species that self-reproduce through autocatalytic cross-reactivity networks (*7, 8*). However, the total sequence variation remains limited to a few nucleotides in substrate-binding regions, which prevents evaluating the impact of autocatalytic self-reproduction in the origin of life.

RNA neutral spaces characterize all the sequences sharing the same property. They have been explored using secondary structure prediction algorithms (*9, 10*), deep mutational scanning (*11*–*19*) and machine learning (*20, 21*). These studies have demonstrated the existence of a connected neutral space in the neighborhood of a reference active sequence. However, the extent of variation studied remains typically limited to within 20 mutations, and none of these studies estimated the size of the neutral space relevant for RNA self-reproduction. Here, using evolutionary conservation in group I introns (Fig. 1A), we devised generative probabilistic models based on statistical learning and structure prediction (Fig. 1B) combined with a high-throughput catalytic assay (Fig. 1CD), which we showed to be a proxy for autocatalysis. This enabled the exploration of a very large neutral space of catalytic RNAs derived from Group I Introns together with estimations of its size.

**Fig. 1.**
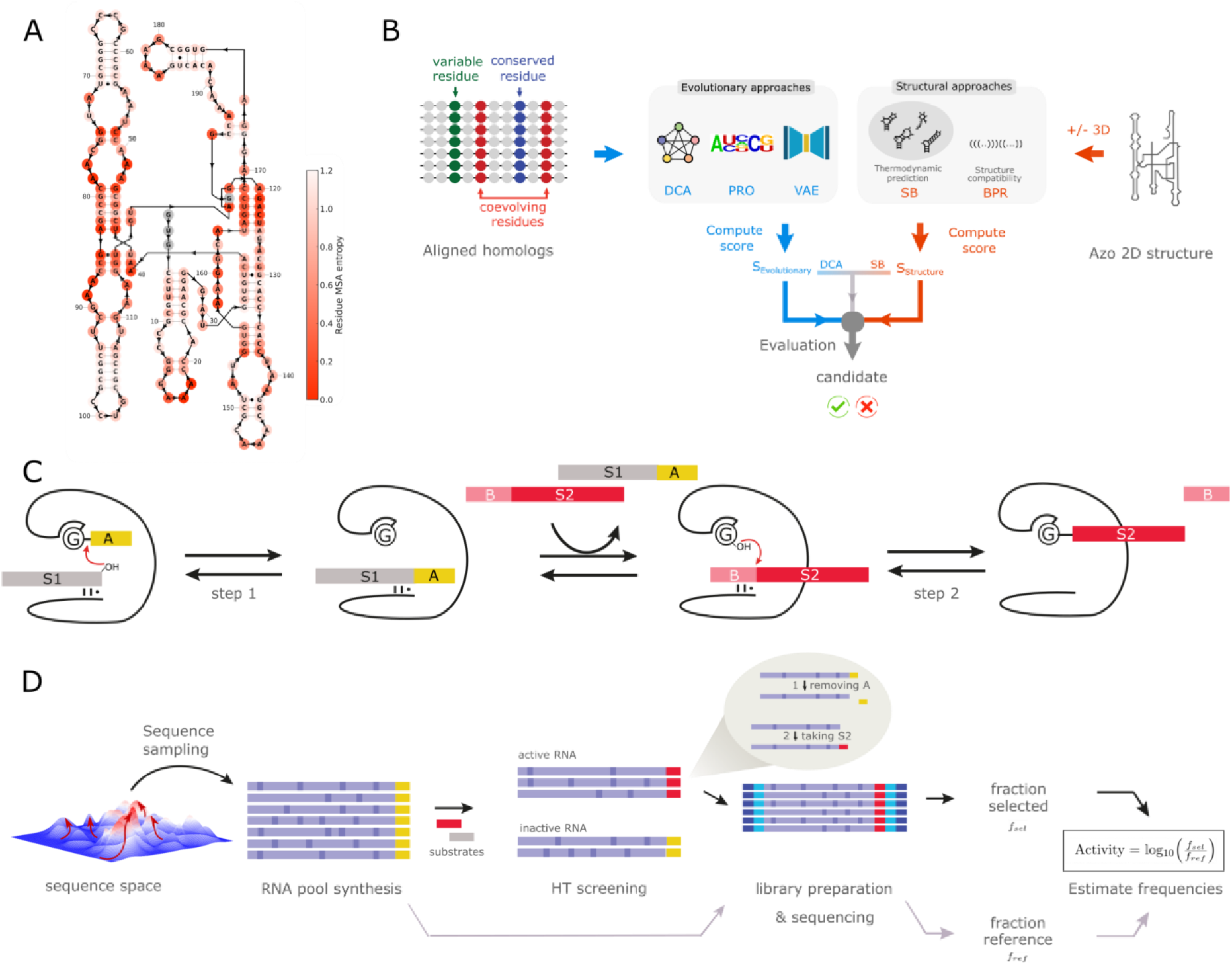
Computational and experimental workflow for the design of artificial group I introns. (**A**) Azoarcus secondary structure. More conserved positions among homologous sequences of the Multiple Sequence Alignment (MSA) appear in darker red. (**B**) RNA were designed using evolutionary models (DCA, VAE, PRO), structural models (SB, SB-3D, BPR), and combination thereof (DCA-SB). The input of evolutionary models is a MSA. The input of structural models is *ab initio* predictions of secondary structure or experimental 3D contacts of Azo. Designs are generated based on probabilistic scores and/or structure scores depending on the model. (**C**) Self-splicing assay: candidate ribozymes (black) are synthesized with an exon sequence A (yellow); RNAs are mixed and incubated with the substrates S1 and B-S2; the depicted two-step reaction transfers the exon A to S2, exchanges the substrates, then ligates S2 onto the ribozyme. (**D**) Screening workflow: after computational generation (panel B), the ribozymes are transcribed from a DNA pool, and tested by the screening assay (panel C); after screening, the active ribozymes are amplified with a substrate complementary primer before being sequenced; the frequency of active sequences (carrying S2) post-assay is normalized by the variant frequency pre-assay.

To assess the generative power of computational models, we developed a deep sequencing assay, with which we analyzed 24,220 unique RNA sequences (Fig. 1CD, Supplementary Material ‘High-throughput self-splicing assay’). This assay mimics the two steps of the natural self-splicing activity of Group I Introns (fig. S1), from which autocatalytic self-reproduction has been engineered (*6, 7*). RNA candidates are first synthesized together with a tRNA fragment exon at their 3’ end (yellow in Fig. 1C). The RNA libraries are then incubated together with two substrate RNAs (gray and red in Fig. 1C). As a proxy for autocatalysis, the RNA is considered active if it transfers its exon to a first substrate, exchanges it for a second substrate, and attaches the extremity of the latter to its own 3’ end (plain red in Fig. 1C). Active RNAs are thus distinguished from inactive ones by deep sequencing, based on the fragment carried at their 3’ end (Fig. 1D). For each variant, we defined the activity score as the logarithm of the fraction of active sequences after screening, divided by the fraction of sequences before incubation, setting the zero at the Azo reference score (Fig. S2). The assay was highly reproducible with a Pearson correlation of 0.99 between independent triplicates (p<10^−5^) (Fig. S3). As the reaction consists of attaching fragments to the catalysts themselves via specific binding to substrates, cross-catalysis is expected to be negligible, which is supported by the absence of activity for a large number of mutations (see below). Additionally, we verified a posteriori that cross-catalysis negligibly affected the activity score by assaying variants separately and as subpools (Fig. S4-S6, Supplementary Methods ‘Cross-catalysis tests’).

We tested several models by evenly populating bins of 10 mutations up to 150 mutations away from Azo (Fig. S2, 150 sequences or more per model per bin). All models displayed a loss of activity as more mutations were introduced in Azo, with the score reaching a lower plateau corresponding to inactive sequences (Fig. S2, Fig. S7). For random uniform mutagenesis (denoted RDM), activity was completely lost after 15 mutations, confirming previous studies (*16*). The measurement noise distribution was taken as the distribution of scores at more than 100 mutations overall all models (Fig. S7), and the threshold for significant activity was set at a z-score of 3.09 corresponding to an activity score of -2.76, and ensuring a p-value<10^−3^ (Supplementary Material ‘Activity Threshold & Active Fraction’). Figure 2A displays the active fraction, defined as the fraction of each variant above the noise threshold, as a function of the number of mutations introduced in Azo. Below, for each generative model, we report L_50_ and L_max_, the largest number of mutations in Azo such that the model generates 50% and 1%, respectively, of active sequences (Fig. 2, Supplementary Table 1, Supplementary Material ‘L_50_ and L_max_’). L_50_ can be interpreted as a typical number of mutations that can be introduced in Azo by a given model, beyond which its predictive power decreases. L_max_ is the maximum number of mutations that we could reliably assess given our experimental resolution per bin. Note that deeper sequencing of larger RNA pools may allow one to assess functional sequences at frequencies below 1%.

**Fig. 2.**
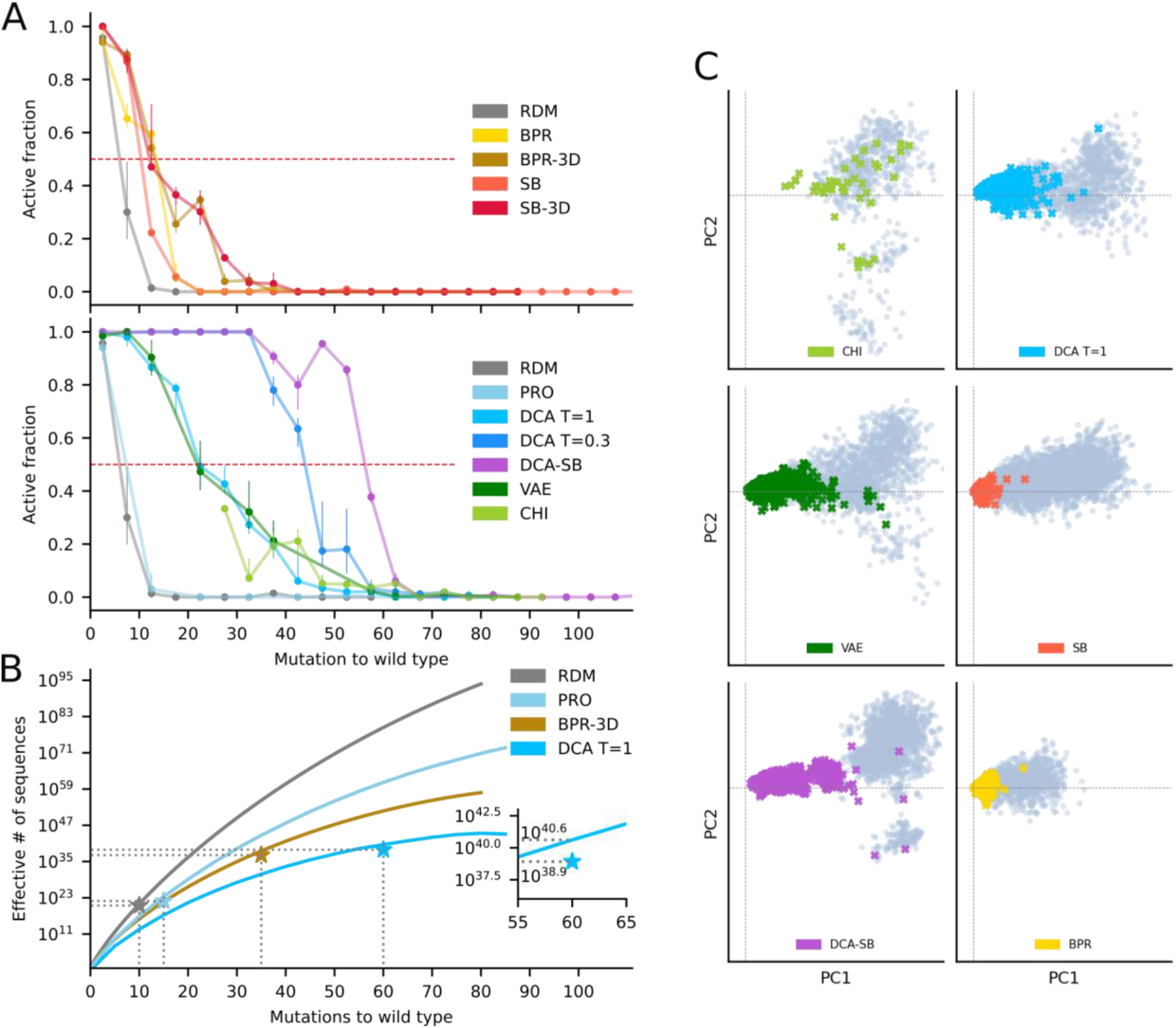
Comparison of the generative power of computational models. (**A**) Fraction of active designs by model as a function of the number of mutations introduced in Azo, by bins of 5 mutations. Dots are the average activity per bin and lines represent the 25-75 percentile range. Top: structure-based models; Bottom: Statistical learning and hybrid models. (**B**) Effective support size of models as a function of mutational distance, showing how many different sequences the model can generate at any given distance. The star indicates the estimation at L_max_. The star values have been corrected with the experimental active fraction at L_max_, which is of order 1% Inset: zoom on the experimental correction at L_max_ for DCA T=1. (**C**) Principal Component Analysis (PCA) of generated sequences (gray) overlayed with active sequences (color). All panels are projected on the same axis system, which are the first two principal components of chimeric sequences (analogs of natural GIIs), centered on Azo, where the first component strongly correlates with the average number of mutations (Pearson ρ=0.88, p<10^−5^).

We first explored the neutral space covered by Group I Introns (GIIs) using Direct Coupling Analysis (DCA), which has been proven to generate *in vivo* neutral variants of protein enzymes (*22*). DCA is a statistical learning approach that accounts for nucleotide conservation and covariation (*23*) via networks of pairwise couplings, and describes a probability distribution over the space of functional nucleotide sequences. Our model was trained on a Multiple Sequence Alignment (MSA) of 815 GIIs sequences that share the same domain composition and secondary structure as Azo (Fig. S8). Although MSA sequences may differ in length due to gaps or insertions relative to Azo, nucleotide covariations can be learned at positions aligned with Azo. Sequences of the same length as Azo (197 nucleotides) were sampled by Markov Chain Monte Carlo up to 90 mutations. DCA generated sequences with L_50_=20 and up to L_max_=60, a large improvement as compared to random mutations, for which L_50_=5 and L_max_=10 (Fig. 2A, table S1). Reaching this degree of mutagenesis using evolutionary conservation was not a priori obvious. Indeed, chimeric sequences (CHI), directly derived from natural GIIs by removing insertions relative to Azo and replacing deletions by the Azo nucleotides (Fig. S9), never reached the 50% success rate per bin (Fig. 2A). Overall, only 6% of them were active, possibly due to the disruption of base-pairs (on average 7 over the 59 base-pairs in the Azo structure).

To estimate the number Ω_DCA_ of potential autocatalytic self-reproducers according to the DCA model, one cannot just correct the total number of possible sequences by the model success rate at a mutational distance. Indeed, DCA biases predictions towards designs to which it attributes high probability. Thus, for a fixed number of mutations, the effective number of sequences among which the model samples is smaller than the total number of possible sequences. Standard results in information theory (*24*) show that the vast majority of the probability mass described by a model is concentrated in a subset of size Ω = exp(S) sequences, where S is the model Shannon entropy −∑_*x*_ *P*(*x*) *logP*(*x*), where the sum is taken over all sequences *x*, and *P*(*x*) is the probability of *x* according to the model. Ω can be interpreted as the effective number of different sequences that the model can generate (the so-called support size of the model). Intuitively, sampling a finite number of sequences from the probability distribution *P*(*x*) is highly likely to be part of this subset of Ω sequences out of the 4^L^ possible sequences. Although it is not known how to compute Ω in general, we devised a semi-analytical method exploiting the structure of DCA (table S2, Supplementary Material ‘Support size computations’) (*25*). This theoretical number increases sub-exponentially with mutational distance, reaching 10^41^ sequences at 60 mutations (Fig. 2B). The success rate at L_max_=60 mutations was determined to be larger than 1% with p < 4.10^−5^ confidence. Thus, correcting Ω_DCA_ with the experimental success rate leads to Ω_DCA_ ≃10^39^.

We further examined the sequence diversity generated by DCA. A Principal Component Analysis (PCA) projection of DCA sequences in the first two principal components of the chimeras (representative of natural diversity) centered on Azo (Fig. 2C) shows that DCA covers the diversity of natural sequences and interpolates with Azo, where PC1 strongly correlates with the distance to Azo (Pearson ρ=0.88, p<10^−5^). To quantify the variety of sequences generated relative to the natural diversity, we computed: For each sequence, its distance from the closest chimera (including Azo), and plotted the distribution of these distances per model (Fig. 3A, top); The distances between all pairs of sequences within each model (Fig. 3A, bottom). Active DCA sequences were found to be on average 25 mutations away from each other (Fig. 3A, bottom), on average 18 mutations away from any chimera, and up to 55 mutations away from any chimera (Fig. 3A, top, 15 Table S3). These distances were comparable to those found in between chimeras: chimeras were on average 32 mutations and up to 59 mutations away from each other (Fig. 3A, Table S3). It thus appears that DCA achieves a form of interpolation, by generating a diversity equivalent to that of natural GIIs (Fig. 3BC), as well as a form of extrapolation, by generating sequences as far away from any chimeras as the latter are from each other. Extrapolation is further confirmed by the largest distance between active DCA designs and chimeras being 99 mutations (Table S3), well beyond the largest distance between any two chimeras.

**Fig. 3.**
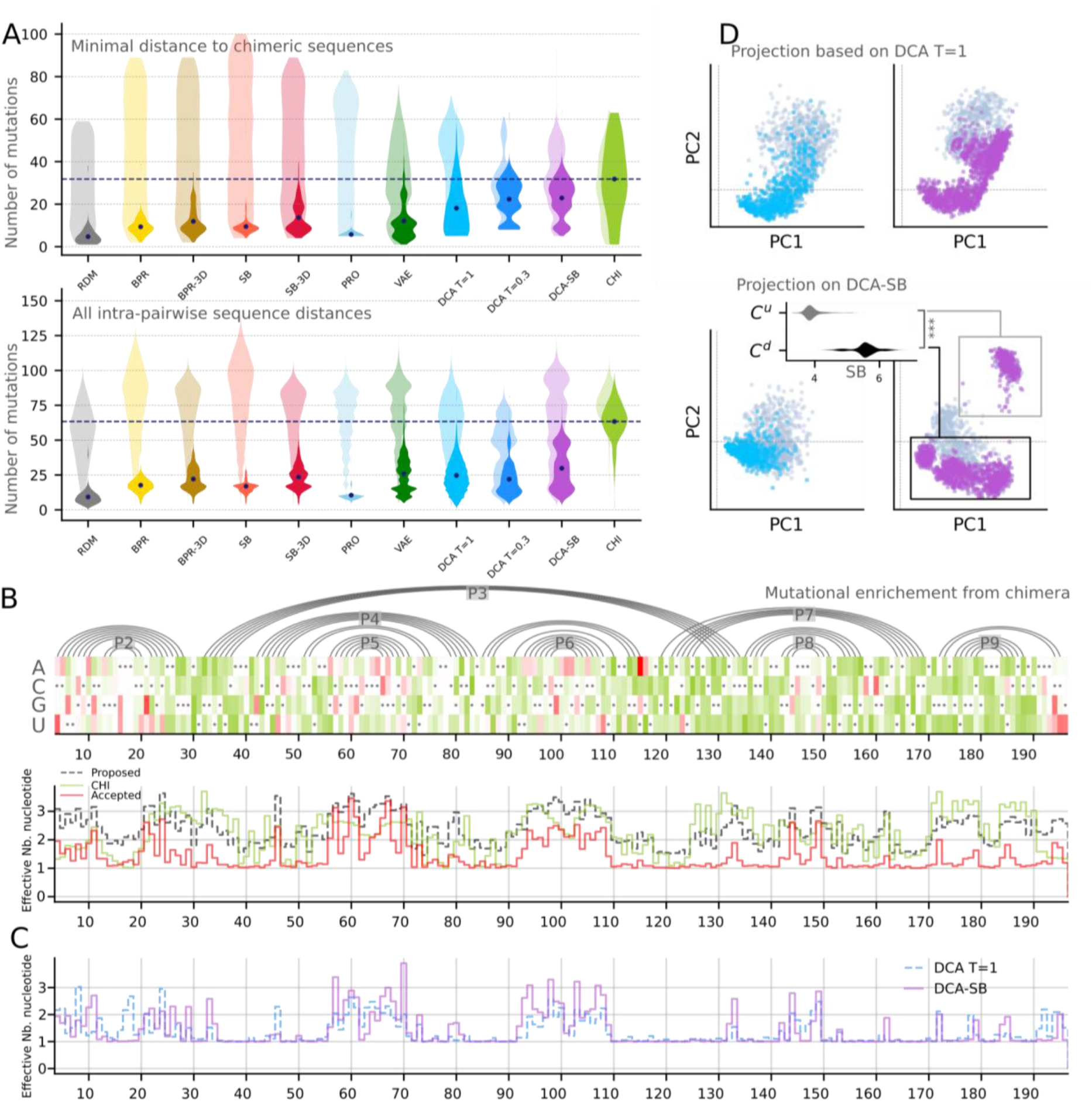
Diversity explored by the generated sequences. (**A**) Upper panel: Violin plots of minimal distance to the set of chimeric sequences for generated sequences (light shades) and active sequences (solid shades), per model ; The average of distances for chimeric sequences is represented by a dashed line. Lower panel: Violin plot of distances between all pairs for generated sequences (light shades) and active sequences (solid shades), per model. **(B)** Mutational enrichment from chimeric sequences. Top: At each position (x-axis), Azo nucleotides are marked by a black dot, shades are the log ratio of mutation frequencies relative to Azo in chimeras (green) and in active variants over all models (red). Bottom: effective number of nucleotides in chimera (green), candidate designs (black) and active designs (red). (**C**) Mutational enrichment due to the structure: Effective number of nucleotides per position in DCA T=1 (blue) and DCA-SB (purple). (**D**) Top: PCA projection of DCA (left) and DCA-SB sequences (right), on the 2 principal components of the DCA T=1 set. Active sequences are colored and inactive ones are gray. Bottom: same but in the principal components of the DCA-SB set, revealing an upper cluster (C^u^) populated by DCA-SB only, distinct from the cluster below (C^d^) that comprises all DCA sequences. Inset: Distribution of structure scores in the two clusters, showing that C^u^ has a significantly better structure score.

Can we explore a larger neutral space? One may seek for more predictive models, able to introduce a larger number of mutations (as quantified by L_50_ and L_max_), or models that yield a larger effective number of generated sequences at a given distance (as quantified by Ω). For instance, random mutagenesis considers all possible mutations and would theoretically yield the highest possible Ω, but its success rate decreases very quickly below our ability to experimentally probe active fractions. This case illustrates the fact that populating the neutral space faces a trade-off. On the one hand, less restricted models may catch a larger number of active sequences but with a lower success rate. On the other hand, more constrained models like DCA generate functional sequences with a higher number of mutations, but at the cost of exploring more limited regions of the sequence space. The outcome in terms of Ω is not obvious, as, for a given model, it depends on how Ω increases with the number of mutations as well as on the L_max_ that can be reached experimentally.

We first tested more relaxed models, which may access a larger diversity but a lesser mutational distance. As discussed, the success rate of random mutagenesis (RDM) decreases dramatically with the number of mutations, with L_50_ = 5 and leading to Ω_RDM_ = 10^20^ at L_max_ = 10, much lower than for DCA (Fig. 2A, Table S1). The profile model (PRO), where each nucleotide is replaced according to its probability in the MSA, is less constrained compared to DCA but retains some evolutionary information. This model is only marginally better than the random one at introducing mutations, with L_50_ also equal to 5 (Fig. 2A, Table S1), but Ω_PRO_=10^22^ at L_max_ = 15. This low number likely results from a high probability of disrupting the secondary structure when selecting nucleotides independently, as 60% of Azo positions are paired (Fig. 1A), as already noted for chimeras. To assess the impact of base-pairing, we randomly replaced base pairs of Azo with other canonical pairs and randomized nucleotides at unpaired positions (base-pair replacement model, BPR). This model performed significantly better than the profile model, with L_50_=15 and Ω_BPR_=10^29^ at L_max_ = 20. Further constraining the model by forbidding variations at tertiary contacts (30 positions in Azo, Fig. S10, BPR-3D model) better populated the tail of the distribution of active RNAs, with L_50_=15 and L_max_=35. Ultimately, Ω_BPR-3D_=10^37^, smaller but close to Ω_DCA_.

We then wondered whether more constrained models managed to introduce a larger number of mutations. Variational AutoEncoders (VAE) directly encode high-order interactions in a neural network, which is relevant to GIIs catalysis as it involves third-order interactions (*26*). Although VAE covered a similar diversity than DCA as shown by PCA projections (Fig. 2C), it performed slightly less well, with L_50_=15 and L_max_=60. This may be explained by the modest number of training sequences, insufficient to resolve couplings of order higher than two (*27*). Alternatively, we introduced constraints by sampling sequences with higher probability according to DCA, which is achieved by lowering the so-called sampling temperature to T=0.3 instead of T=1 for the standard version. This indeed allowed for the introduction of more mutations, with L_50_=45 and L_max_=60.

As hinted by the base-pair replacement model (BPR), structural constraints can help introduce more mutations while preserving catalysis. To account for the global structure, we generated sequences based on thermodynamic secondary structure prediction (SB model for ‘structure-based’) (*28*), using a heuristic that retained the pseudo-knot as a probabilistic signature in the contact map (Fig. S11) (*29*). However, this model led to a performance close to that of random mutations, with L_50_=10 and L_max_=20. This poor performance could be due to discrepancies between the predicted and the actual structure (Fig. S11) or to structure energy minimization being non-optimal for catalysis (Fig. S12). Fixing positions involved in 3D contacts (SB-3D model) strongly improved performance, with L_50_=10 and L_max_=40, significantly beyond mere base pair replacement. This shows that SB models account for more cooperative effects than BPR, as expected, but only when critical tertiary interactions are unperturbed.

We finally combined the statistical and structural approaches in the DCA-SB model, including secondary structure and any other constraint imposed during evolution. This allowed for the largest number of successful mutations, with L_50_=55 and L_max_ = 65, with a significantly higher success rate in the range 60-70 compared to DCA alone (two-sample t-test, p < 10^−5^). Interestingly, the DCA-SB sequences introduced diversity in the loops P5 and P6 (Fig. 3C, 1A), a diversity which was not found in DCA alone. This is confirmed by PCA projections, where the DCA-SB sequences covered the DCA cluster plus an additional cluster characterized by a better structure score (Fig. 3D).

For the models able to accumulate more mutations than the standard DCA, namely DCA T=0.3 and DCA-SB, we could compute their support size only over a limited range of mutational distances (Table S2). Nevertheless, over the explored range of mutations, both models were found to populate a much more restricted fraction of the sequence space, with Ω_DCA T=0.3_ and Ω_DCA-SB_ being many decades below Ω_DCA_ (Table S2).

As the largest estimation for the number of active ribozymes thus arises from the standard DCA model, we confirmed on the DCA pool that the assay is a proxy for self-reproduction for the generated sequences by assaying individual sequences at varying distances from Azo up to 60 mutations (Supplementary Material ‘Self-reproduction assay’). We tested autocatalysis from two and four substrate fragments (Fig. 4, Fig. S13) (*6*). More than 60% of the sequences (N=17) with an assay score larger than the activity threshold were found to reproduce from fragments, up to 60 mutations.

**Fig. 4.**
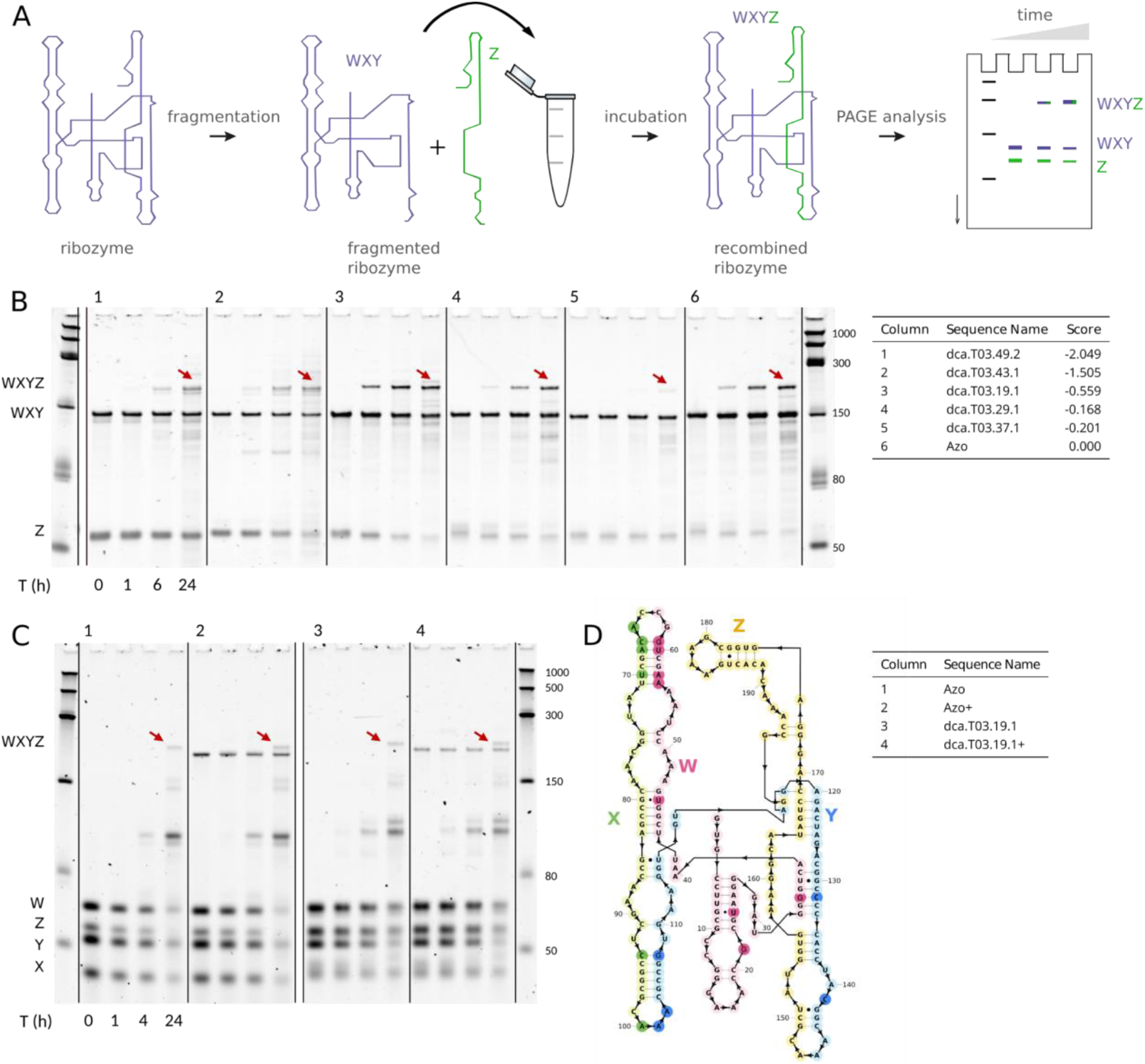
Evidence for self-reproduction. (**A**) Two-fragment self-reproduction experimental assay. The ribozymes are first fragmented into two pieces (WXY and Z) at position 145, then incubated at defined time points. The formation of longer covalent copies is monitored by gel electrophoresis. (**B**) Two-fragment self-reproduction assay for 5 designs and Azo, assayed for 0, 1, 6 and 24 hours. The assayed ribozymes are documented in the table on the right of the gel: the number of mutations from Azo is indicated in their name, and their activity score for the self-splicing assay in the last column. The red arrows are pointing at the recombined covalent ribozymes. For other designed ribozymes with 15 to 60 mutations, see Fig. S13. (**C**) Four-fragment self-reproduction assay for one design and Azo, assayed for 1, 2, 4 and 24 hours. The dca.T03.19.1 design and Azo are first fragmented into four fragments (W, X, Y and Z) at the positions 62, 99 and 145. Two conditions were assayed : the four fragments in absence of the covalent ribozyme; and the four fragments in presence of the covalent ribozyme (condition +). The red arrow shows the presence of the fully recombined covalent ribozymes. (**D**) The dca.T03.19.1 ribozyme. The colors represent the 4 different fragments, and the nucleotides colored with darker shade highlight the position of mutations compared to the wild type Azo.

Until now, only a handful of autocatalytic RNA self-reproducers had been proven to exist, which did not allow any meaningful assessment of plausibility given the 10^118^ possible sequences of the length of Azo (to be compared with 10^81^ estimated atoms in the observable universe). Based on the comparison of several statistical models and the consistency of their predictive power over thousands of measurements each, the DCA model provided the largest lower bound for the number of RNA self-reproducers, which is of order 10^39^. Although this estimation does not yet prove that an RNA emergence of life is probable, it now sets the problem within the typical scales of physics and chemistry (Supplementary Material ‘Plausibility of RNA self-reproduction’). In any case, it indicates that abiogenesis could have followed many alternative paths. Importantly, this estimation is only a lower bound. Indeed, ligase ribozymes were selected from random pools of size 10^15^ (*30*), suggesting that generative models capable of successfully populating larger neutral spaces should exist. The existence of a larger neutral space is already supported by the fact that we could diversify a GII other than Azo (Fig. S14, Supplementary Material ‘Phormidium’), and the existence of structurally distinct autocatalytic RNAs (*31*). The density of self-reproducers may thus be many decades higher, making evidence for a probable RNA origin of life within reach.

## Acknowledgments

The authors acknowledge Vincent Messow and Nono S. C. Merleau for preliminary work, Sandeep Ameta and Michal Matyjasik for discussions.

## Funding

Institut Pierre-Gilles de Gennes ANR-10-EQPX-34 (CL, VO, PN)

EU H2020 Grant ERC AbioEvo 101002075 (CL, VO, PN)

Human Frontier Science Program RGY0077 / 2019 (CL, VO, PN)

EU H2020 grant MSCA-RISE In-ferNet 734439 (MW)

## Author contributions

Conceptualization: EH, MW, MS, PN

Experimental Methodology: CL, EH, PN

Computational Methodology: VO, FC, FZ, MW, MS

Investigation: CL, VO, FC

Funding acquisition: EH, MW, MS, PN

Project administration: PN

Supervision: FZ, EH, MW, MS, PN

Writing – original draft: CL, VO, PN

Writing – review & editing: CL, VO, FC, FZ, EH, MW, MS, PN

## Supplementary Materials

Materials and Methods

Supplementary Text

Figs. S1 to S18

Tables S1 to S4

Data and programs are available at:https://zenodo.org/records/13138337

## References and Notes

1. W. Gilbert, Origin of life: The RNA world. nature 319, 618–618 (1986).

2. J. Attwater, A. Raguram, A. S. Morgunov, E. Gianni, P. Holliger, Ribozyme-catalysed RNA synthesis using triplet building blocks. Elife 7, 35255 (2018).

3. N. Papastavrou, D. P. Horning, G. F. Joyce, RNA-catalyzed evolution of catalytic RNA. Proceedings of the National Academy of Sciences 121, 2321592121 (2024).

4. P. Pavlinova, C. N. Lambert, C. Malaterre, P. Nghe, Abiogenesis through gradual evolution of autocatalysis into template-based replication. FEBS letters 597, 344–379 (2023).

5. T. A. Lincoln, G. F. Joyce, Self-sustained replication of an RNA enzyme. Science 323, 1229–1232 (2009).

6. E. J. Hayden, N. Lehman, Self-assembly of a group I intron from inactive oligonucleotide fragments. Chemistry & biology 13, 909–918 (2006).

7. N. Vaidya, M. L. Manapat, I. A. Chen, R. Xulvi-Brunet, E. J. Hayden, N. Lehman, Spontaneous network formation among cooperative RNA replicators. Nature 491, 72–77 (2012).

8. J. T. Sczepanski, G. F. Joyce, Synthetic evolving systems that implement a user-specified genetic code of arbitrary design. Chemistry & biology 19, 1324–1332 (2012).

9. P. Schuster, W. Fontana, P. F. Stadler, I. L. Hofacker, From sequences to shapes and back: a case study in RNA secondary structures. Proceedings of the Royal Society of London. Series B: Biological Sciences 255, 279–284 (1994).

10. W. Fontana, P. Schuster, Shaping space: The possible and the attainable in RNA genotype-phenotype mapping. Journal of Theoretical Biology 194, 491–515 (1998).

11. J. I. Jiménez, R. Xulvi-Brunet, G. W. Campbell, R. Turk-MacLeod, I. A. Chen, Comprehensive experimental fitness landscape and evolutionary network for small RNA. Proceedings of the National Academy of Sciences of the United States of America 110, 14984–14989 (2013).

12. S. Kobori, Y. Yokobayashi, High-Throughput Mutational Analysis of a Twister Ribozyme. Angewandte Chemie - International Edition 55, 10354–10357 (2016).

13. O. Puchta, B. Cseke, H. Czaja, D. Tollervey, G. Sanguinetti, G. Kudla, Molecular evolution: Network of epistatic interactions within a yeast snoRNA. Science 352, 840–844 (2016).

14. C. Li, W. Qian, C. J. Maclean, J. Zhang, The fitness landscape of a tRNA gene. Science 352, 837–840 (2016).

15. J. Domingo, G. Diss, B. Lehner, Pairwise and higher-order genetic interactions during the evolution of a tRNA. Nature 558, 117–121 (2018).

16. E. J. Hayden, D. P. Bendixsen, A. Wagner, Intramolecular phenotypic capacitance in a modular RNA molecule. Proceedings of the National Academy of Sciences 112, 12444–12449 (2015).

17. A. D. Pressman, Z. Liu, E. Janzen, C. Blanco, U. F. Müller, G. F. Joyce, I. A. Chen, Mapping a systematic ribozyme fitness landscape reveals a frustrated evolutionary network for self-aminoacylating RNA. Journal of the American Chemical Society 141, 6213–6223 (2019).

18. J. O. Andreasson, A. Savinov, S. M. Block, W. J. Greenleaf, Comprehensive sequence-to-function mapping of cofactor-dependent RNA catalysis in the glmS ribozyme. Nature communications 11, 1663 (2020).

19. J. M. Roberts, J. D. Beck, T. B. Pollock, D. P. Bendixsen, E. J. Hayden, RNA Sequence to Structure Analysis from Comprehensive Pairwise Mutagenesis of Multiple Self-Cleaving Ribozymes. eLife 12, 80360 (2023).

20. R. Rotrattanadumrong, Y. Yokobayashi, Experimental exploration of a ribozyme neutral network using evolutionary algorithm and deep learning. Nature communications 13, 4847 (2022).

21. S. Sumi, M. Hamada, H. Saito, Deep generative design of RNA family sequences. Nature Methods 21, 435–443 (2024).

22. W. P. Russ, M. Figliuzzi, C. Stocker, P. Barrat-Charlaix, M. Socolich, P. Kast, R. Ranganathan, An evolution-based model for designing chorismate mutase enzymes. Science 369, 440–445 (2020).

23. F. Cuturello, G. Tiana, G. Bussi, Assessing the accuracy of direct-coupling analysis for RNA contact prediction. RNA 26, 637–647 (2020).

24. T. M. Cover, J. A. Thomas, Elements of Information Theory (Wiley, New York, 1999).

25. F. Calvanese, C. N. Lambert, P. Nghe, F. Zamponi, M. Weigt, Towards parsimonious generative modeling of RNA families. Nucleic Acids Research 52, 5465–5477 (2024).

26. S. Li, M. Z. Palo, X. Zhang, G. Pintilie, K. Zhang, Snapshots of the second-step self-splicing of Tetrahymena ribozyme revealed by cryo-EM. Nature Communications 14, 1294 (2023).

27. F. J. Poelwijk, M. Socolich, R. Ranganathan, Learning the pattern of epistasis linking genotype and phenotype in a protein. Nature communications 10, 4213 (2019).

28. R. Lorenz, S. H. Bernhart, C. Siederdissen, H. Tafer, C. Flamm, P. F. Stadler, I. L. Hofacker, ViennaRNA Package 2.0. Algorithms for molecular biology. 6, 1–14 (2011).

29. J. S. McCaskill, The equilibrium partition function and base pair binding probabilities for RNA secondary structure. Biopolymers: Original Research on Biomolecules 29, 1105–1119 (1990).

30. D. P. Bartel, J. W. Szostak, Isolation of new ribozymes from a large pool of random sequences. Science 261, 1411–1418 (1993).

31. R. Mizuuchi, N. Ichihashi, Minimal RNA self-reproduction discovered from a random pool of oligomers. Chem. Sci. 14, 7656–7664 (2023).

