## Supplementary Materials for "Expanding the space of self-reproducing ribozymes using probabilistic generative models"

**The PDF file includes:**

Materials and Methods
Supplementary Text
Figs. S1 to S18
Tables S1 to S4

### Materials and Methods

High-throughput self-splicing assay

The assay aims at discriminating against an activity similar to self-splicing, the reaction catalyzed by the wild type Azoarcus GII. Figure S1 shows the two steps of self-splicing reaction, where the GII is flanked by two exons denoted A and B, one on each side. For the natural self-splicing reaction, the GII binds a free GTP, allowing the attack of A. Then, the 3' of A attacks the 3' of the GII to allow the formation of a covalent bond between the two exons, while releasing the GII.

To mimic this self-splicing mechanism, we devised an experimental assay comprising two steps, as shown in Figure 1C. All ribozymes were produced with a 15-nt long sequence at their 3' extremity, called sequence A, which corresponds to the wild type exon of Azoarcus GII. First, the sequence A is transferred at the end of the substrate called S1. Then, a second substrate B-S2 binds into the recognition site of the RNA, leading to the transfer of S2 at the 3' end of the ribozyme and the release of the B fragment. The ribozymes with the sequence S2, thus considered active, were selected during the later steps.

We performed three large pooled experiments, wherein we assayed thousands of sequences simultaneously. For each computationally designed pool of RNA sequences, we ordered the corresponding DNA templates with the exon at the 3' end, and the T7 promoter in the 5' end. The single strand DNA oligo pools containing 12 000 or 18 000 sequences were purchased from Twist Bioscience. We amplified the DNA pools by PCR (~15 cycles) with the KAPA Hifi HotStart ReadyMix (Roche), and purified the samples with the NucleoSpin Gel and PCR cleanup (Macherey Nagel). The DNA templates were then transcribed using the HiScribe T7 High Yield RNA Synthesis Kit (New England Biolabs) for 4 hours at 37°C to produce RNA molecules. The samples were then subjected to phenol-chloroform extraction and ethanol precipitation using 0.1 volume of 3M Sodium Acetate (Sigma) and 2.5 volumes of cold 100% ethanol. After extraction, the samples were treated with the DNAse I (New England Biolabs) and PAGE purified on a 8% urea PAGE.

For each designed RNA pool, we performed two sub-experiments. The first sub-experiment is the self-splicing assay. The second one is a control experiment to correct for the biases of relative quantity of each synthesized ribozyme within the corresponding pool (before reaction), used to compute the activity scores. For the self-splicing assay, 2 $\mu M$ of ribozymes were incubated with 25 $\mu M$ of substrates S1 and B-S2 in a buffer (30 mM EPPS pH7.5, 60 mM $MgCl2$) at 37°C in a final volume of 20 $\mu L$. Two samples were taken during the incubation, at 0 and 60 mins, mixed with loading solution (70% formamide, 130 mM EDTA, 0.1% xylene cyanol, 0.1% bromophenol blue) and loaded on a 8% Urea PAGE. The reaction was quenched by adding 60 mM of EDTA, and the ribozymes cleaned using the Monarch RNA cleanup kit (New England Biolabs) with an adjusted volume of ethanol and binding buffer. For the control experiment, no substrate was used and the ribozyme pool was directly cleaned with the Monarch RNA cleanup kit.

The RNA samples were then prepared for sequencing, using the NEBNext Ultra II Directional RNA Library Prep Kit for Illumina (New England Biolabs). For the self-splicing experiment, the primer used during the reverse-transcription corresponds to the sequence complementary to the S2 substrate, whereas the sequence complementary to the sequence A was used for the control experiment. The samples were sequenced on a NovaSeq SP flow cell (2∗250 nts, 2∗800 M reads) in paired ends and with 25% of PhiX by the NGS platform at Institut du Cerveau et de la Moelle épinière (ICM, Paris).

To test the robustness of our experimental protocol across the 3 pools, we introduced a common set of 355 sequences, which showed a very consistent activity computation across all pools ($\rho = 0.99$ correlation between each pair of pools, Fig. S3).

Cross-catalysis tests

Cross-catalysis within a pool would result in false positives (apparently active ribozymes that are in reality inactive), which in turn could lead to overestimation of the lower bound on the number of ribozymes. To test for this, we performed a series of experiments ensuring that cross-catalysis was negligible.

First, we assayed 24 sequences by denaturing gel electrophoresis, individually (Fig. S4). We used identical reaction conditions as the pooled high-throughput assay: 2 $\mu M$ of ribozymes incubated with 25 $\mu M$ substrates at 37°C during 1h. These sequences were selected among the pool designed by DCA to cover the range of mutations 15-60 from Azo, with various scores of the assay as measured by sequencing. Overall, all the sequences that had a sequencing score above the threshold also displayed a form of catalytic activity visible on the gel. 1 in the 24 sequences was a false negative : it displayed products on the gel but a sequencing score below the threshold. However, neither false negatives impact our lower bound estimation of the number of ribozymes, nor they reveal cross-catalysis as the latter results in false positives (Fig. S5).

Second, we measured by the sequencing assay the activity score of a set of sequences spanning a range of activity scores, comparing the score obtained in a pool and as separate individual measurements. The sequences were produced, purified, and tested separately. The scores reproduced well the relative experimental activity measured in the pool, with $\rho$=0.95 with p-value < $10^{-5}$ (Fig. S6).

We further tested that active ribozymes did not affect the measured activity of each other within a pool as a function of their relative concentration, by taking a subset of 12 only active sequences, tested as a subpool. The activity score of this subset incubated as a separate smaller pool correlated strongly with the scores within the total pool, with $\rho$=0.98 with p-value < $10^{-5}$ (Fig. S6), showing that the overall concentration of active variants did not affect the activity score.

Self-reproduction assay

The fragmentation strategy used to fragment the Azoarcus ribozyme (*1*) was generalized to the artificial ribozymes. We fragmented the ribozymes at the nucleotide positions 145-147, which we referred to as the Y/Z junction. This fragmentation site is located in the loop of the P8 paired region Fig. 4.A). For each ribozyme, two fragments were produced with the addition of short sequence fragments necessary for the recombination reaction to occur. For the first fragment (WXY fragment), a 'CAU' tag was inserted in 3' and the second fragment (Z fragment), a 'GGCAU' tag was inserted in 5' by PCR.

The self-reproduction assays were carried out as follows. 1 $\mu M$ of each ribozyme fragments were incubated in a buffer (30 mM EPPS pH7.5, 60 mM MgCl2) at 37°C. Samples were taken at different time points during the incubation, mixed with loading solution (70% formamide, 130 mM EDTA, 0.1% xylene cyanol, 0.1% bromophenol blue) and loaded on a 8% Urea PAGE. The gels were dyed with SyBr Gold (ThermoFisher) diluted in TBE buffer.

Furthermore, we fragmented one ribozyme into 4 fragments (W, X, Y Z fragments), according to (*1*) (Fig. 4D). The 'CAU' tag was inserted in 3' of the fragments W, X and Y and the 'GGCAU' tag was inserted in 5' of the fragments X, Y and Z. Similarly, the self-reproduction reaction was performed by incubating 1 $\mu M$ of each 4 fragments in the incubation buffer (30 mM EPPS pH7.5, 60 mM MgCl2) at 37°C. The course of the reaction was followed by analyzing samples on a 8% Urea PAGE.

### Supplementary Text

Construction of the MSA

To build the MSA, we used the wild type sequence (197 nucleotides long) of Azoarcus GII and its known secondary structure derived from the pdb X-ray structure 1G9B (*2*) in order to detect homologous sequences from the RFAM database (*3*). To account for similarity but also secondary structure compatibility, we built a seed alignment containing only the Azoarcus sequence and its secondary structure, which we fed to the alignment algorithm implemented in the package Infernal (version 1.1.4) (*4*) to search for homologs in the 2611 sequences of the RF00028 RFAM family. We filtered out all the sequences for which the computed e-value is > $10^{-3}$, which is the probability of finding the obtained alignment score randomly. 816 sequences were found below the homology threshold of $10^{-3}$ and aligned using Infernal. Figure SI.4 shows the diversity per position in this alignment, measured as the exponential of Shannon entropy, and the propension of gaps per nucleotide position.

Calculation of activities from sequencing results

To estimate the experimental activities, we computed the frequencies of designed sequences in: i) the reference condition before the catalytic reaction, and ii) the reacted condition where the substrate has been mixed with the designs then incubated. For both conditions, we mapped each paired-end read to the closest designed sequence using the software Blast (version 2.12) (*5*). Then, we selected the reads that covered at least 70% of the mapped designed sequence with full identity, which allowed us to obtain the occurrence of each design in the sample. We computed the frequencies $f_{ref}$ of designs in each pool before catalysis, allowing us to quantify the bias in RNA molecules in the initial synthetized pool. Then, we computed the frequencies of designs $f_{sel}$ in the sample with the substrate. To do so, we counted the reads when the substrate was attached right after the 3' end of the design, which indicates that the ribozyme was able to excise the exon and then ligate the substrate. Finally, we computed the experimental activity $act = log\left(\frac{f_{ref}}{f_{sel}}\right)$. Analyzing reverse reads was sufficient for calculating the activity score. For the ambiguous cases where designs that counted less than 5 reads in the pool before catalysis were excluded from the analysis. Sequences that had $f_{ref}$ > 0 and $f_{sel}$ = 0 were deemed as inactive.

Activity threshold & active fraction

As depicted in Figure S2, as more mutations are introduced, the activity score decreases from zero (the Azo score reference) to a lower plateau. This plateau is reached for all models beyond 70 mutations, characteristic of non-functional sequences. We considered the distribution of scores at this plateau representative of the experimental noise, taking the score distribution over the 543 sequences comprising at least 100 mutations (red in Fig. S7). The noise distribution is found to be Gaussian (Fig. S7). Significant activity was taken for a p-value < $10^{-3}$, resulting in a threshold of -2.76. Note that this p-value guarantees a z-score larger than 3.09 (3 standard deviations above the mean). For reference, activities greater than -2.45 corresponded to p-value<$10^{-4}$, and activities greater than -2.16 to p-values<<$10^{-5}$. Those more stringent thresholds are verified for most of the active sequences found in the $L_{max}$ bin, and the presence of multiple sequences above the threshold further lowers the level of evidence, thus lowering the p-value (see sections below and Table S1). For each model, we computed the active fraction per bin spanning 5 mutations of distance from Azo. For each bin, the active fraction is defined as the number of designs with activity levels above the threshold divided by the total number of designs in that bin.

The error bars on the bins active fraction are estimated using the error on the individual sequence activity measurements. For this, we used data from 355 overlapping sequences, whose activity has been independently measured three times. As already pointed out, these measurements displayed high consistency, with a correlation coefficient close to 0.99 across different experiments. For each of these 355 sequences, we calculated the standard deviation of the activity measurements. The vast majority of the standard deviations are below 0.3 (with just 5/355 designs exceeding this value). Consequently, we considered 0.3 as the error on the single activity measurement. Using this error estimate, we computed the error bars for the active fraction in each bin by counting the number of designs whose activity values crossed the threshold, considering the ±0.3 measurement error.

### L50 and Lmax

$L_{50}$ was estimated as the largest number of mutations such that a given model achieved at least 50% success rate within a bin of 5 mutations with p-value<$10^{-3}$ (binomial test, Table S1). $L_{max}$ was estimated as the largest number of mutations such that a given model achieved at least 1% success rate within a bin of 5 mutations with p-value<$10^{-3}$ (binomial test, Table S1).

However, as the $L_{max}$ bin generally comprises more than 1 sequence, and with scores clearly above the threshold, we computed and reported the p-value given all the available data, rejecting the hypothesis that all sequence activities in this bin are drawn from the noise, as explained below and reported in Table S1. To compute this actual p-value for $L_{max}$, we assumed a Gaussian null-model for the measured activities $a$ of non-functional sequences, $a \sim N(\mu, \sigma)$, which we call the noise model. As is described above in Section "Activity threshold & active fraction", the model mean $\mu$ and standard deviation $\sigma$ are estimated from a sample of 543 non-functional sequences having at least 100 mutations. We assume here that all measured activity is due to experimental noise.

Assume now a sample of $N$ sequences $\{s_i, i = 1, \ldots, N\}$ with experimentally measured activities $\{a_i, i = 1, \ldots, N\}$ (typically the sequences of any bin in mutational distance to Azo). For any arbitrary $z$-value, we can find the number $n(z) = |\{i, a_i > \mu + z\sigma\}|$ of sequences with measured activity being at least $z$ standard deviations above the mean of the noise model. We can also calculate the probability $\pi(z) = P(a > \mu + z\sigma)$ that a randomly chosen activity $a \sim N(\mu, \sigma)$ from the noise model is beyond that activity threshold.

These two numbers, via a one-sided binomial test, give access to a $z$-dependent p-value (which allows to refuse or not the hypothesis that all data are drawn from the null model) :

$$p = \sum_{i=n(z)}^{N} \frac{N!}{i!\,(N-i)!}\,\pi(z)^i\,[1-\pi(z)]^{N-i}$$

In the paper, in coherence with our selection threshold chosen according to Section "Activity threshold & active fraction", we report p-values for sequences above our standard threshold, i.e. for $\pi(z) = 0.001$, reached at. $z \simeq 3.09$.

In principle, any $z$-value could be used for calculating $p$-values. For smaller $z$, we would have more super-threshold activities (larger $n(z)$) but of smaller individual significance (smaller $\pi(z)$), while a larger $z$ would lead to less selected sequences of higher individual significance. Note that fixing our activity threshold (and thus considering one value of $z$) leads to an upper bound of the $p$-values.

### Models

The distribution of nucleotide frequencies for each model is provided as Figure S15.

#### *Profile (PRO)*

Each position is drawn independently from the distribution of nucleotides at the corresponding position in the MSA. In this case, the frequencies are computed while omitting gaps to sample full sequences. The distribution of nucleotides per position is shown in Figure 3B and Figure S15.

#### *Base-Pair Replacement (BPR)*

The complementary design strategy consists in sampling only sequences that are compatible with the known secondary structure of Azoarcus GII. Note that this is only a compatibility condition, which does not guarantee that the minimum free energy structure is indeed the Azoarcus GII. In practice, for paired positions in the structure, including the pseudoknot, we sampled pairs of nucleotides that can form canonical base pairs (A, U), (C, G), and (G, U). For the unpaired position, nucleotides were drawn uniformly. Therefore, sequences sampled with this strategy are compatible with Azo structure, which however does not mean the minimum free energy structure is that of Azo.

#### *Direct Coupling Analysis (DCA)*

In Direct Coupling Analysis, each nucleotide sequence is described by a Boltzmann-like probabilistic model: $P(n1 \dots nL) \propto exp\,(-H(n1, \dots nL))$, where the Hamiltonian $H(n1, \dots nL) = \sum h_i(a) + \sum J_{ij}(a, b)$ represents the log-likelihood. Here, high probability sequences are sequences that restitute best the statistical signature of natural counterparts.

The training of the DCA T=1 model was conducted using the methods and code provided in (*6*). Sequences were sampled from the model at fixed distances from the Azoarcus reference sequence, with distance values ranging from 5 to 90 in increments of 5. For each distance value, 150 sequences were sampled using Gibbs sampling, ensuring equilibrium was reached. The generation was biased toward the reference sequence using a biasing potential $H(n1, \dots, nL) =$

$\sum hi(a) + \sum Jij(a,b) + \theta \cdot distance(a, azo)$. For each distance value, the 150 sequences were randomly selected from those below the 10% quantile of the corresponding distance DCA energy (10% best DCA score). The training of the DCA model for DCA T=0.3 and DCA+sb was conducted using the procedure described in (*7*). To determine the parameter $h_i$ and $J_{ij}$, we started with the observed frequencies $f(n,i)$ of nucleotide $n$ at each position $i$ in the MSA.

The parameters were updated in order to reproduce these single and pairwise frequencies (Fig. S16). For the learning procedure, we used N=20 × $10^3$ MC sweeps, K=20 × $10^3$ MC steps. MC sweeps were run at kT=0.3 with a regularization λ = 0.3. Once the model trained, we compared the predicted frequencies $f_i, f_{ij}$ to the ones observed in MSA and obtained Pearson's correlation of $\rho = 0.99$ for $f_{ij}$, and $\rho = 0.96$ for connected correlations $c_{ij}(a,b) = f_{ij}(a,b) - f_i(a) \times f_j(b)$, the latter being a consistency check are they not directly fitted by the procedure (Fig. S16). This update has been done using a non-persistent contrastive divergence algorithm as described in (*7*), see pseudo-code in Figure S17.

We generated sequences by sampling first $k$ positions to be mutated, which were drawn randomly from 10 to 100 by step of 10 mutations. For each $k$ positions selected, we performed 100 sweeps of $k \times 200$ MCMC steps used in the metropolis test with the DCA energy parametrized above. Then, we pooled all the designed sequences and populated bins of 10 (from 0-10 to 80-90) with the 280 sequences with the best DCA score and at least 5 mutations from each other.

*Variational AutoEncoder (VAE)*

For the variational autoencoder (VAE), we derived our implementation from reference Ding et al. (*8*). A VAE is composed of three main elements: an encoder neural network, a decoder neural network, and a latent space. The encoder (denoted He) is used to find the underlying data structure of the MSA by projecting sequences into a low dimensional latent space $Z$. The main difference of this latent space is that $Z$ is modeled as a Gaussian variable.

To generate sequences, the decoder network is used to convert latent representation into sequences. To train this model, we used the evidence lower bound (ELBO) loss, which composed of two terms:

$$ELBO(\theta,\phi) = \sum_{Z} q_\phi(Z \mid X) \, log \, p_\phi(X \mid Z) - \sum_{Z} q_\phi(Z \mid X) \, log \, \frac{q_\phi(Z \mid X)}{p_\phi(Z)}$$

where the θ are the weights of the encoder and the φ are the ones for the decoder. For each sequence, this loss evaluates first the log-likelihood predicted by the decoder network (the first term), then compute the divergence of the $Z$ representation produced by the encoder from its prior Gaussian distribution (the second term).

To choose the best architecture, we tried several hyper parameters with a 5-fold cross-validation procedure. We selected the architecture with two layers of 512 hidden units (for both encoder and decoder), and 128 dimensions for the latent space, with a ReLU activation function. The decoder predicts log-probabilities that are converted back using the Softmax function over the four nucleotides. To read the predicted sequence from the output of the decoder, we chose at each position the nucleotide with the highest probability.

To sample artificial RNA sequences, we drew randomly 60000 data points in the latent space using a Gaussian distribution centered at the coordinate of the wild type using several variances $\sigma \in 0.1, 0.9, 1, 10$. Then, we decoded all latent data points into sequences. Then, we populated bins of 10 with 280 randomly selected sequences, from 1 to 100 mutations, where at least five mutations are observed between each pair of designs.

#### *Structure-Based (SB)*

To predict the secondary structure of a sequence, we used the thermodynamic energy model called nearest neighbor with the Turner2004 parameters (*9*). Here, the parameters gives the free energy of folding of a sequence into a given structure, where the associated probability is $p(s) = exp\{-\Delta G(s)\}$. The Zucker algorithm (*10*), a dynamic programming algorithm, allowed us to compute the minimum free energy structure of a given sequence for this energy model. However, this algorithm does not account for pseudoknots.

To measure the selectivity of one sequence to adopt a specific secondary structure among all possible secondary structures, we used McCaskill's algorithm (*11*), which is a variant of Zucker's algorithm. This algorithm enumerates all possible secondary structures, in contrast with the prediction of the minimum free energy structure, and records the probability of each pair of positions to be paired in this ensemble. The probabilities of pairing are reported into a matrix called base pair probability matrix (BPPM). We obtained the BPPM using the implementation of McCaskill's algorithm in the ViennaRNA package (version 2.5.17) (Fig. S11).

Similarly to Zucker's algorithm, this does not account for pseudoknots; however, we noticed residual probabilities at positions involved in the pseudoknot, as shown in Figure S11. We compared the BPPM predicted by ViennaRNA, and the algorithm of NUPACK (*12*) that explicitly accounts for pseudoknots. The prediction with accounting explicitly for pseudoknots does not yield a drastic improvement while being one order of magnitude slower.

The structure recovery score (SB) is based on the difference between the predicted BPPM and the known secondary structure of Azoarcus GII, where $\delta_{ij\in\sigma}$ is 1 if positions $ij$ are paired in the known structure (denoted $\sigma$ here). $BPPM_{i,j}$ is the predicted probability of $i, j$ are paired for the sequence $s$.

#### *Tertiary structure constraints (3D)*

To favor the recovery of tertiary interactions and the catalytic core, we devised additional constraints where we fixed the wild type nucleotide of Azoarcus at positions involved in the IGS, the terminal G, the tetraloop motif (GAAA), the P7-loop involved in the catalytic core, and positions involved in tertiary interactions as delineated in (*13*) using an X-ray structure and molecular dynamics simulation. In total, 66 out of 197 positions were held fixed during the sequence space exploration, as shown in Figure S10. We imposed these constraints in the context of the BPR model and the SB model.

*Direct Coupling Analysis & Secondary structure (DCA-SB)*

To generate sequences from DCA and the secondary structure score, we combined both scores in the metropolis test of the MCMC with an acceptance score $exp\left((\alpha DCA(s) + SB(s))/T\right)$. To sample sequences, we performed the same protocol as for the DCA alone generated sequences, considering several $\alpha$ values: $0.1, 0.3, 0.4, 0.7$. The goal was to reduce the importance of the DCA contribution, enabling the exploration of sequence driven by the biophysical model. Once the sequences were sampled, we pooled all the designs and populated bins of 10 mutations (from 0-10 to 90-100 mutations) with the 280 sequences having the best DCA score, but with at least 5 mutations from each other.

*Experimental benchmarking*

We benchmarked the DCA scores and the SB score on experimentally generated data published earlier (*14*){Citation}. In this study, the authors started with the wild type sequence of Azoarcus GII, and synthesized a pool of variants where five positions were randomly mutated. To ensure sufficient diversity, they also randomly introduced additional mutations using PCR prone mutagenesis. Using this protocol to generate sequence diversity, they designed four pools of many thousand RNA sequences, representing four strengths of selection pressure for the catalytic activity. To control the strength of the selection pressure, the authors used the concentration of MgCl2 (mM), allowing them to discriminate the very active variants that were able to perform the selected catalytic activity with low MgCl2 ('Str' at [MgCl2]=2 nM), see Table S4. In contrast, poorly active variants only appeared in the non-selected batch ('Pre').

To test our scoring functions (DCA and SB), we first aligned each sequence to the wild type one, using the global alignment algorithm that only accounts for matches and mismatches. Because of the experimental protocol used in this reference, 20 positions were not mutated. Figure S18 shows the distribution of DCA scores (relative to the wild type, which is set to zero) per pool of mutants, where lower is better. As expected, the distribution is ordered by the strength of the selection pressure which has been used; therefore, showing that the DCA score indeed captures to some extent the catalysis of GII. Moreover, the lowest DCA score here is zero (the wild type), consistent with the assumption that natural GII are more efficient than artificial mutants. Similarly, in Figure S18, we performed the same calculation with the SB score, which also yielded comparable results. We compared the SB score with the ensemble defect (ED) (*12*) where both performed similarly. We chose the SB score because it displayed a larger difference between the *Pre* (no selection) and *Str* (strongest selection) pools.

Contact predictions

To predict the contacts between pairs of positions using $H^{azo}$, we computed the Average Product Correction (APC) $F_{ij}^{APC}$ for each pair of positions. For this system, we used the 1% highest $F_{ij}^{APC}$ (188 pairs of positions) as prediction of contacts (Fig. S11). We compared the prediction with the X-ray structure (contacts are defined by a cutoff of 3.5Å on the minimum distance between nucleotides heavy atoms). 54.7 % of the predictions are correct according to the secondary structure or the considered X-ray structure, while 83% of the secondary structure contacts were correctly predicted. Moreover, some tertiary contacts were also recovered (dark circles, Fig. S11).

Phormidium

To confirm our results, we designed sequences starting not from Azoarcus wild type GII but Phormidium GII—a 208 nucleotide long sequence. For Phormidium, we built the MSA based on the 2611 sequences of GII in RFAM (RF00028), where we selected 1424 sequences (compared to the 815 selected for Azoarcus), which we then aligned. To search and align Phormidium homologs, we used Infernal with a seed constituted only with Phormidium wild type sequence obtained from the database GISSD and a secondary structure obtained by Shape-Map experiment (Fig. S14).

From the DCA model, we designed 1341 sequences across 10 bins of 10 mutations (0-10, 10-20, …) populated with roughly 140 sequences each. Similarly with Azoarcus, we combined DCA with the SB score, where the DCA score is scaled with $\alpha = 0.5$. To sample sequences with mutations ranging from 0 to 90, we used seven MCMC sweeps of $2.\dot{1}0^6$ steps with a biasing potential based on the distance to the wild type to drive the MCMC toward more or less mutated sequences. Figure S14 shows the activity measured by deep sequencing across the different bins of number mutations. These results support our findings with Azoarcus: i) Azoarcus is not a special point as our design exploration also works with another GII, and ii) the SB score improved the exploration of the sequence space by a similar amount. DCA alone displayed an $L_{50}$=25, whereas the contribution of secondary structure yielded an $L_{50}$=30.

Support size computations

The support size of a probability distribution is the space of all outcomes that have a non-zero probability. In contrast, the effective support size is an estimation of the number of different outcomes you can expect when sampling from the distribution. These two concepts are different. For instance, with a fair dice, you expect 6 possible different outcomes. However, for a rigged dice where the face with the number 6 has a probability of $1/1000$ (compared to the standard $1/6$), you would expect something closer to 5 different outcomes when sampling from it.

The effective support size is measured as the exponential of the probability distribution Shannon entropy. For a uniform probability distribution, the effective support size and the support size are identical (fair dice effective support size: $2^{2.58\dots} = 6$). For a non-uniform one, it better represents the expected different outcomes (rigged dice effective support size: $2^{2.33\dots} = 5.03\dots$). In information theoretic terms (*15*), the effective support size can be defined as the minimal number of sequences collecting the dominant majority of the probability mass described by the probability distribution.

At any given distance (number $K$ of mutations) from the wild type, the sequences we designed with our models are drawn from the probability distribution $P_{model}(n|K)$. At each distance, we can compute the effective support size for each model as the exponential of the entropy of the probability distribution. Not all generated sequences will be functional; by testing the designs, we estimate the success rate as the fraction of sequences that are active. The product of the effective support size and the success rates is consequently an estimate of the potential number of functional sequences at a given mutational distance.

For example, randomly introducing 10 mutations from the wild type results in approximately $10^{21}$ possible sequences, cf. below for details. At that distance, for the random designs, we observe an active fraction of 0.3 for our designs. This means that potentially, 30% of the random $10^{21}$ mutated sequences are functional, leading to an estimate of around $10^{20}$ potentially working sequences.

For some of our control models (Random, Random Base Pairs, Random Base Pairs + 3D), since each possible outcome is equiprobable, the support size $\Omega_K$ coincides with the effective support size. The estimation for these models is done using combinatorial calculations of the number of sequences fulfilling the model constraints at a distance of $K$ mutations from Azo.

*Random Mutations*

Calculating the support size for random mutations consists of determining how many possible mutations exist for the wild type sequence a given distance $K$. To do this, we count in how many different ways we can choose $K$ sites out of 193 and consider that each site can mutate in 3 different ways (since there are 3 possible mutations per site if we exclude the original base).

$$\Omega_K = \frac{193}{K} \cdot 3^K$$

*Random Base Pairs*

To compute the support size for the secondary structure-compatible random mutations, consider the 59 Watson-Crick contacts (including those involved in the pseudoknot) and 75 'free' residues not engaged in Watson-Crick pairing. Each of the 75 free residues can mutate in 3 different ways. All the Watson-Crick pairs can mutate in 5 different ways (considering also the wobble pair GU).

The mutational possibilities for the wild type pairs are diagrammed as follows:

| Original pair | 2 mutations from the wild type | 1 mutation from the wild type |
|---|---|---|
| AU | UA, CG, GC, UG | GU |
| GC | UA CG, AU, UG | GU |
| GU | UA, CG, UG | GC, AU |

Except for the wobble pairs GU, a base pair can undergo 4 types of mutations when both residues mutate, and 1 type of mutation if only one residue mutates. Since only 6/59 pairs are GU in the wild type, the calculation has been done assuming the behavior of Watson Crick pairs for all secondary-structure pairs.

We define $z$ as the number of Watson-Crick pairs mutating by introducing one mutation relative to the wild type, $y$ as the number of pairs introducing two mutations from the wt, and $x$ as the number of unpaired sites mutating. The combinatorial size for $K$ mutations can be expressed as:

$$\Omega_K = \sum_{z=1}^{59} \frac{59}{z} \sum_{x+2y=K-z} \frac{75}{x} \cdot 3^x \cdot \frac{59-z}{y} \cdot 4^y$$

where the second sum is performed over the solution of the equation considering that

$$x \in \{0, \cdots, 75\}$$
$$y \in \{0, \cdots, 59-z\}$$

For any given $z$ in the first sum, the condition on the second sum ensures that we reach exactly $K$ mutations from the wild type.

*Random Base Pairs 3D*

The calculation of the support size for the RBP-3D is exactly the same as for the RBP, except that 17 Watson-Crick pairs and 28 free residues are kept fixed. We are left with 59-17 = 42 Watson-Crick pairs and 75-28 = 47 free residues

$$\Omega_K = \sum_{z=1}^{42} \frac{42}{z} \sum_{x+2y=K-z} \frac{47}{x} \cdot 3^x \cdot \frac{42-z}{y} \cdot 4^y$$

with:

$$x \in \{0, \cdots, 47\}$$
$$y \in \{0, \cdots, 42-z\}.$$

*Energy based models*

For the energy based models such as Profile, DCA T=1, covariance, DCA+SB, and DCA T=0.3, each outcome at a given distance is not equiprobable, so we need to estimate the effective support size as the exponential of the Shannon entropy.

The model's energy function $H(n)$ defines a probability distribution over the sequence space, $P(n) = 1/Z \ exp\ (-H(n))$ . To estimate the entropy of $P(n \mid K)$, we use Bayes' theorem, which connects these two probabilities. Indicating with $B_K$ the set of sequences at distance $K$ from the wild type we have:

$$P(n|K) = \frac{P(n,K)}{P(K)} = \frac{P(n) \cdot \delta\{K, dist(n,wt)\}}{P(K)} = \frac{1}{Z} \frac{exp\ \{-H(n)\} \cdot \delta\{K, dist(n,wt)\}}{P(K)}$$
$$S_K = log\ (Z) + [H(n)]_{B_K} + log\ (P(K))$$

where $P(K)$ is the probability that a generated sequence is at distance $K$ from the wildtype (wt). Analytically it is the sum of $P(n)$ over all sequences at that distance,

$$P(K) = \sum_{n \in B_K} exp\{-H(n)\}$$

but it can be computed by generating a number of sequences and counting how many of them are at distance $K$. The partition function $Z$ is unknown and requires a summation over the entire sequence space, which is impossible to perform in practice.

Adding a biasing potential to the model $H_\theta(n) = H(n) + \theta \cdot dist(n, wt)$, as done for the sampling of DCA T=1 designs, does not change $P(n|K)$ or $S_K$ :

$$P_\theta(n|K) = \frac{\frac{1}{Z_\theta} exp\{-H_\theta(n)\} \cdot \delta\{K, dist(n, wt)\}}{\frac{1}{Z_\theta} \sum_{n \in B_K} exp\{-H_\theta(n)\}} = \frac{\frac{1}{Z} exp\{-H(n)\} \cdot \delta\{K, dist(n, wt)\}}{\frac{1}{Z} \sum_{n \in B_K} exp\{-H(n)\}}$$
$$= P(n|K)$$

We use this property, along with the fact that $S_0 = 0$ (the mutational space at 0 mutations only has the wild type, so the probability distribution $P(n|0)$ has only one possible outcome and an entropy $S_0 = 0$ ) to compute $S_K$ for all $K$ of interest.

We take a series of biasing potentials $\{\theta_1, \theta_2, \ldots, \theta_N\}$ such that $\theta_1$ provides good coverage of the mutational distances in $\{0, \ldots, K_1\}$ (meaning that sampling from $H(n) + \theta_1 \cdot dist(n, wt)$ we generate enough sequences with the specified mutational distances to allow for statistical analysis). Similarly $\theta_2$ ensures good coverage from $\{K_1, \ldots, K_2\}$, and in general, $\theta_i$ provides good coverage in $\{K_{i-1}, \ldots, K_i\}$.

The biasing potential $\theta_1$ is strong enough that some of the generated sequences will be at distance 0 from the wildtype. Using the entropy formula at distance 0, we obtain $log(Z_{\theta_1})$

$$S_0 = log(Z_{\theta_1}) + [H_{\theta_1}(n)]_{B_0} + log(P_{\theta_1}(K = 0))$$
$$log(Z_{\theta_1}) = -log(P_{\theta_1}(K = 0)) - [H_{\theta_1}(n)]_{B_0}$$

Having the value of $log(Z_{\theta_1})$, we can compute $S_K$ for all distances $\{0, \ldots, K_1\}$ for which we have a large enough sample to reliably estimate $P_{\theta_1}(K)$ and $[H_{\theta_1}(n)]_{B_K}$. We then relax the bias to $\theta_2$ to increase the distance from the wt. To compute $log(Z_{\theta_2})$, we cannot again use the $S_0 = 0$ trick since the less biased model $H_{\theta_2}(n)$ will not generate enough sequences at distance 0 from the wt. However we know that there is $K^1$ with good overlap between the $\theta_1$-biased model and the $\theta_2$-biased model, so we can compute $S_{K^1}$ from the $\theta_1$ model and then use it to compute $log(Z_{\theta_2})$ .

$$S_{K^1} = log(Z_{\theta_1}) + [H_{\theta_1}(n)]_{B_{K^1}} + log(P_{\theta_1}(K^1))$$
$$S_{K^1} = log(Z_{\theta_2}) + [H_{\theta_2}(n)]_{B_{K^1}} + log(P_{\theta_2}(K^1))$$
$$log(Z_{\theta_2}) = S_{K^1} - log(P_{\theta_2}(K^1)) - [H_{\theta_2}(n)]_{B_{K^1}}$$

This procedure can be iterated to compute $log(Z_i)$ for all the biased models and $S_K$ for all desired distances, considering that:

$$S_{K^{i-1}} = log\ (Z_{\theta_{i-1}}) + [H_{\theta_{i-1}}(n)]_{B_{K^{i-1}}} + log\ (P_{\theta_{i-1}}(K^{i-1}))$$
$$S_{K^{i-1}} = log\ (Z_{\theta_i}) + [H_{\theta_i}(n)]_{B_{K^{i-1}}} + log\ (P_{\theta_i}(K^{i-1}))$$
$$log\ (Z_{\theta_i}) = S_{K^{i-1}} - log\ (P_{\theta_i}(K^{i-1})) - [H_{\theta_i}(n)]_{B_{K^{i-1}}}$$

As observed in table S3, for DCA T=0.3 and DCA-SB the support size estimation was not possible after a certain number of mutations due to ergodicity problems. This means that the Markov Chain Monte Carlo (MCMC) simulations do not fully thermalize and we cannot estimate equilibrium properties like the entropy. However, this does not hinder sequence generation since the chains, despite not being thermalized, still explore regions with favorable model scores (low energy). DCA T=1 support sizes have been reduced by a factor of ten because we only experimentally tested the sequences in the top 10% quantile of the DCA score. This approach is conservative since it assumes that the remaining 90% of sequences are non-active.

Plausibility of RNA self-reproduction

Based on DCA corrected by the experimental success rate, we estimated that there are at least $0{,}02 \times 10^{40{,}62} > 8.10^{38}$ autocatalytic RNAs among $4.10^{118}$ possible sequences of length 197 nucleotides, resulting in a frequency of $f$=$2.10^{-80}$. Let us consider typical scales relevant to astrobiology:

- $2.10^{23}$ stars in the universe
- $10^{21}$ liters of water in an Earth-like ocean
- $3.10^{11}$ attempts to generate sequences during a billion year given an RNA lifetime of 24 hours
- $6.10^{23}$ molecules per liter (1 molar)

If every star would have an Earth-like planet, multiplying these numbers leads to $n$=$4.10^{79}$ RNAs generated overall. As $n \cdot f \approx 1$, a self-reproducing RNA may be typically found over a billion years. Obviously, there is not one such planet per star, it is not possible to expect a molar of RNA, in particular not of length 197 nucleotides. However, this lower bound estimation is now within the scales discussed in physics and chemistry. Beforehand, considering 1 in $10^{118}$ sequences would not have allowed any meaningful assessment. Improving the estimation of the present work by several decades may ultimately lead to a plausible emergence of self-reproduction from random pools of RNA.

**Fig. S1 to S18.**

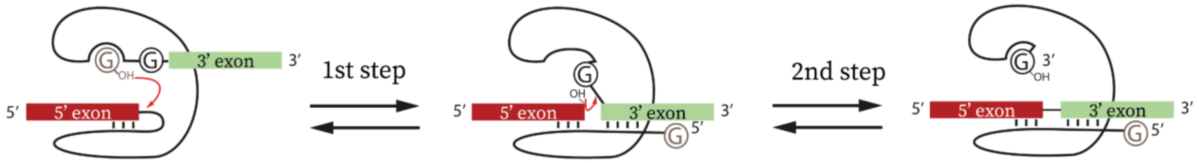


**Fig.1. Self-splicing reaction scheme catalyzed by natural GII**. The self-splicing is a two-step reaction leading to the excision of the ribozyme (shown as a black line) and the ligation of the two exons surrounding the GII (in red and green). The first step involves a free GTP bound in the catalytic pocket (in brown) which attacks the bound between the 5' exon placed at the IGS site of the ribozyme, and lead to the release of the 5'exon and the ligation of the G at the 5' of the GII. The second step of splicing starts by a conformation change of the GII which positions the ending G of the GII (black G) into the catalytic pocket and the 3' exon in close proximity to the 5' exon. The second attack is performed by the last nucleotide of the 5'exon, typically a U involved in a conserved Wobble pair with the IGS of the GII, leading to the breakage of the bound between the ribozyme and the 3' exon and the ligation of the two exons together.

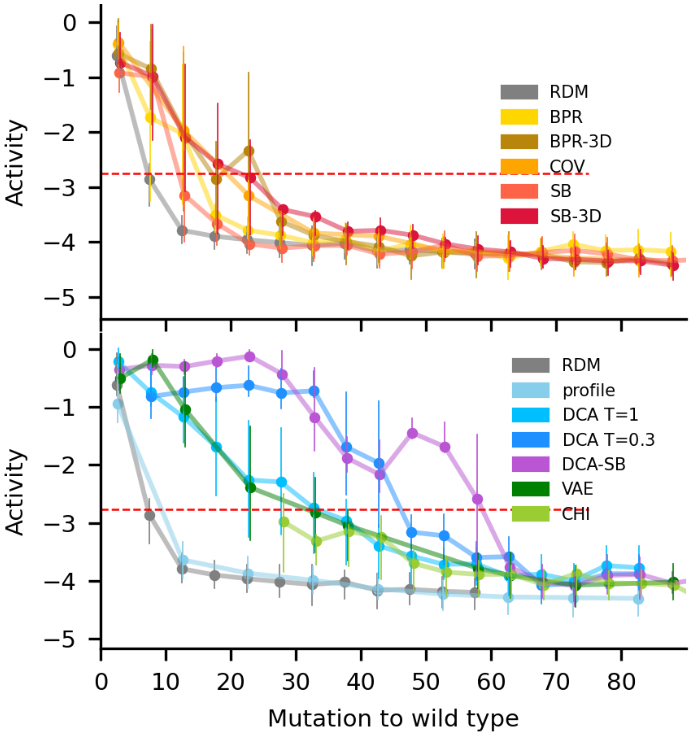


**Fig. S2. Experimental activity as a function of the number of mutations.** Top: biophysics-based generative methods; Bottom: statistical-based generative methods. Both graphs report the experimental activity computed from the sequencing experiments binned every 5 mutations. The dots are the mean activity of each bin while the vertical lines are the first and third quartile. The red dashed line is the active threshold set at a z-score of 3.09 or equivalently a p-value=$10^{-3}$, which corresponds to an activity of -2.76.

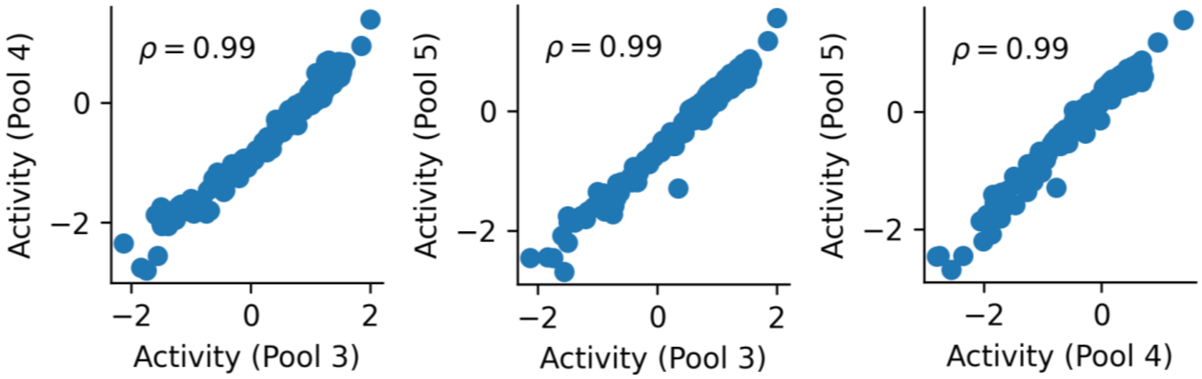


**Fig. S3. Correlation of the activity scores between independent triplicates.** We compared the activity computed for the same set of sequences from 3 independent experiments called Pool 3, Pool 4, and Pool 5. We reported the Pearson correlation ρ for each comparison ($N=355$, $p < 10^{-5}$).

A

Inactive

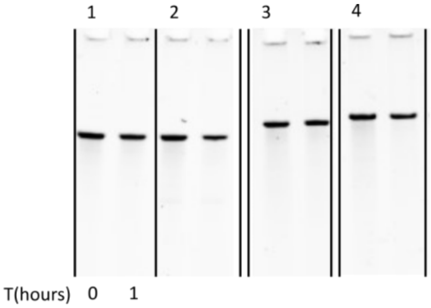


| Column | Sequence Name | Step 2 | Step 1+2 |
|---|---|---|---|
| 1 | dca.T1.30.1 | -2.275 | -4.698 |
| 2 | dca.T1.45.1 | -1.964 | -4.687 |
| 3 | dca.T1.60.3 | -1.450 | -3.046 |
| 4 | dca.T1.45.2 | -1.804 | -3.440 |

B

Canonically Active

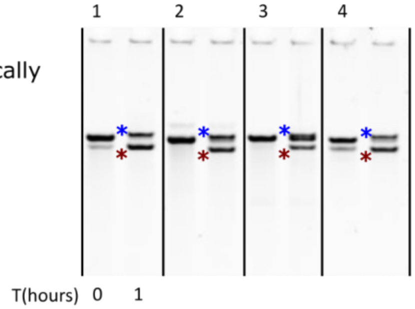


| Column | Sequence Name | Step 2 | Step 1+2 |
|---|---|---|---|
| 1 | dca.T1.15.4 | 0.511 | 0.479 |
| 2 | dca.T1.30.5 | -0.375 | -0.397 |
| 3 | dca.T1.45.5 | -0.321 | -0.357 |
| 4 | Azo | 0 | 0 |

C

Active

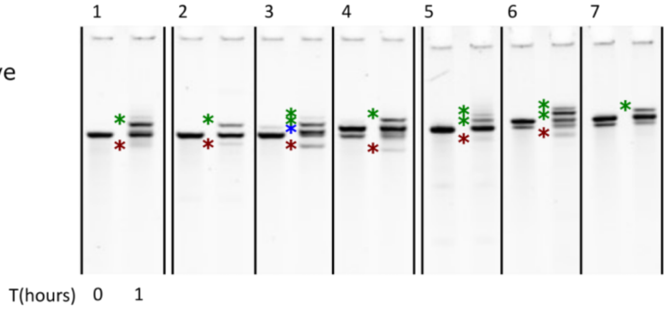


| Column | Sequence Name | Step 2 | Step 1+2 |
|---|---|---|---|
| 1 | dca.T1.15.3 | -0.156 | -1.059 |
| 2 | dca.T1.45.3 | -0.788 | -2.168 |
| 3 | dca.T1.30.4 | 0.215 | -1.499 |
| 4 | dca.T1.45.4 | -0.075 | -1.168 |
| 5 | dca.T1.55.1 | -0.358 | -1.522 |
| 6 | dca.T1.60.1 | 0.570 | -1.232 |
| 7 | dca.T1.60.2 | 0.007 | -1.330 |

D

Only step 2

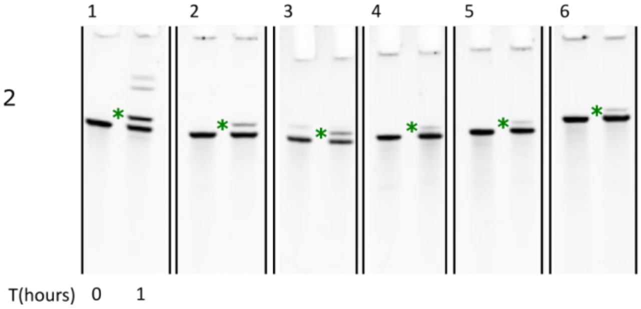


| Column | Sequence Name | Step 2 | Step 1+2 |
|---|---|---|---|
| 1 | dca.T1.15.1 | -0.164 | -4.047 |
| 2 | dca.T1.30.3 | -0.713 | -2.498 |
| 3 | dca.T1.45.6 | -0.599 | -1.365 |
| 4 | dca.T1.50.1 | -0.612 | -1.293 |
| 5 | dca.T1.50.2 | -1.110 | -1.654 |
| 6 | dca.T1.60.4 | -1.109 | -2.870 |

**Fig. S4. Gel electrophoresis analysis of 24 DCA RNAs self-splicing assay**. Left: Denaturing gel electrophoresis of self-splicing assay, grouped by category. Red stars indicate products of step 1, blue stars products of step 1 followed by step 2, green stars products of step 2 without step 1. Inactive: only the candidate ribozyme is visible, none of the products are visible. Canonically active: products of step 1 and step 2 are visible, such that step 1 mostly occurs before step 2. Active: products of step 1 and step 2 are visible, but step 2 may happen while step 1 has not yet occurred. Only step 2: only products of step 2 are visible. Right: scores as measured in the pooled sequencing assay, for step 2 only and steps 1 followed by step 2 (denoted step 1+2). See Figure S5 for the analysis of the correspondence between categories as seen on gels and by sequencing scores.

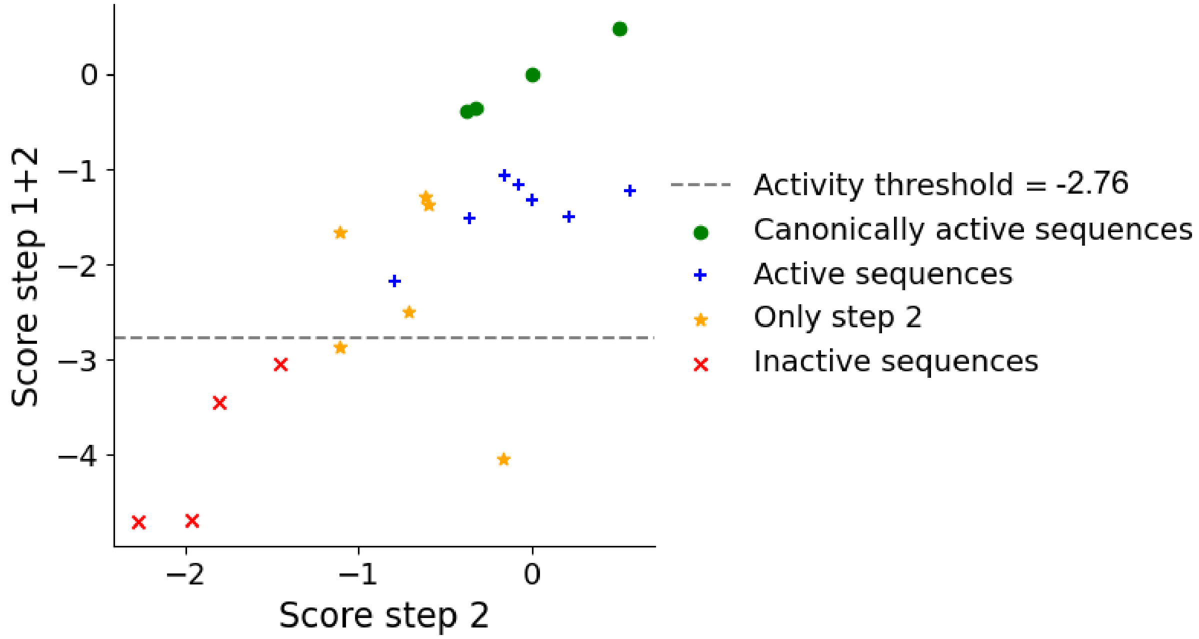


**Fig. S5. Comparison of sequencing scores and products visible by gel electrophoresis.** Categories as seen from electrophoresis gels of Figure S4, as a function of sequencing scores for step 2 only and combined scores (step 1+2). The activity threshold is set at -2.76 on the y-axis (step 1+2). All designs above this threshold indeed lead to visible products (green, blue, yellow). One yellow design is slightly below the threshold, and one is clearly below, indicating false negatives (activity visible on gel but not in sequencing). However, pooled assay false negatives neither reveal cross-catalysis nor impact our lower bound estimation of the number of ribozymes. Canonically active designs from gels (step 1 band and step2 after step 1 band) score highest on both scores. Active designs from gels (displaying step 1 and step 2 products performed in any order) have intermediate step 1+2 score and large step 2 score consistent with the possibility for step 2 to occur first. Designs only showing step 2 on gels have intermediate values for both scores and include a false negative (bottom right yellow dot).

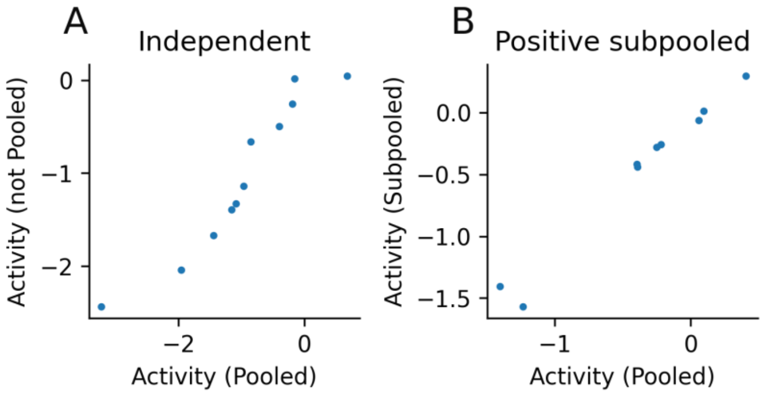


**Fig. S6: Effect of pooling on activity scores.** Comparison of sequencing-based scores: (**A**) For a subset of sequences spanning a range of activity scores, as measured within a large pool of 18 000 sequences (x-axis) versus measured separately from each other; (**B**) Subset of sequences positive in the large pool assay (x-axis, together with 18.000 sequences), versus activity score of the same subset measured when incubated as a subpool.

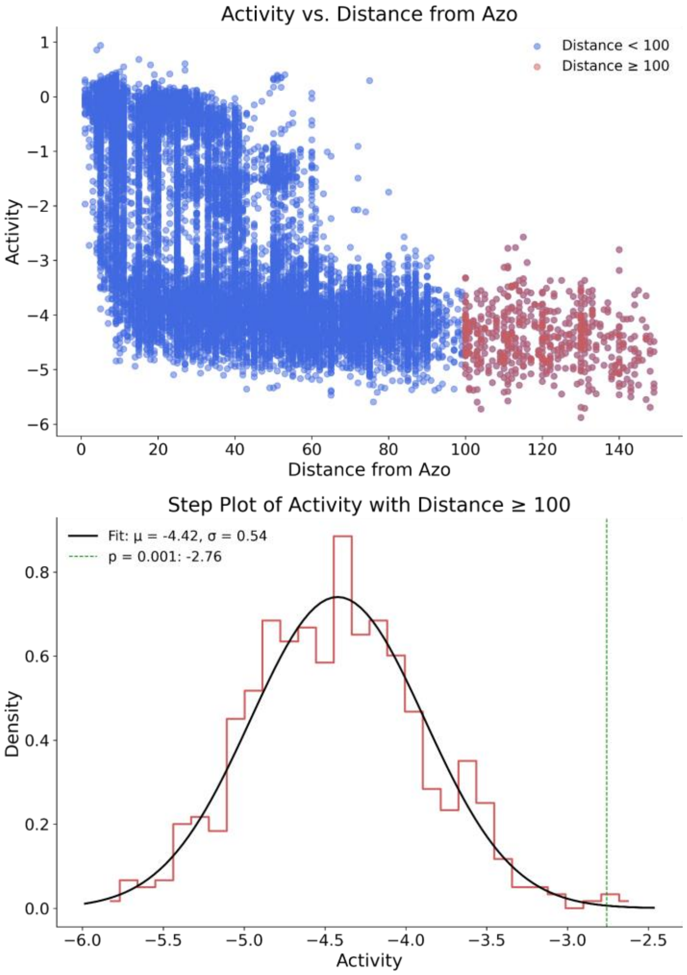


**Fig. S7. Activity score noise distribution.** Top: Scatter plot of the experimental activity against the distance from the wild type for the 15147 sequences that exhibited experimental activity ($f_{sel} > 0$). In red, the 543 designs that introduce 100 or more mutations on the Azo reference. Their activity values are considered representative of experimental noise. Bottom: Gaussian fit on the activities of the 543 designs with 100 or more mutations, used to model the experimental noise. Kolmogorov-Smirnov (K-S) test $p = 0.943$, indicating a 94.3% probability that the observed sample could come from the reference normal distribution (Gaussianity rejection: $p < 0.05$). The vertical line corresponds to the noise threshold of $p = 0.001$.

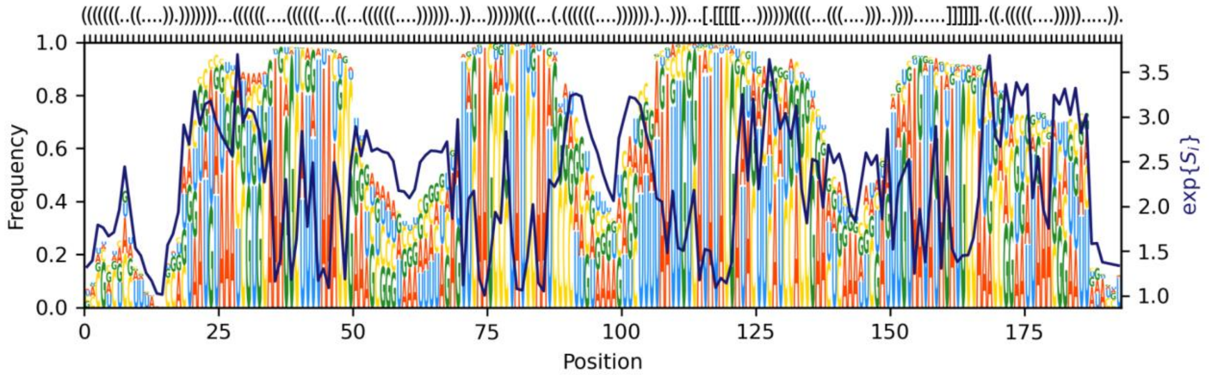


**Fig. S8. Natural diversity by position in the MSA.** Top row: Secondary structure of Azo in dot bracket notation. Main panel: Logo representation indicating the frequency of nucleotides per position. The solid curve represents the effective number of nucleotides per position (exponential of Shannon entropy).

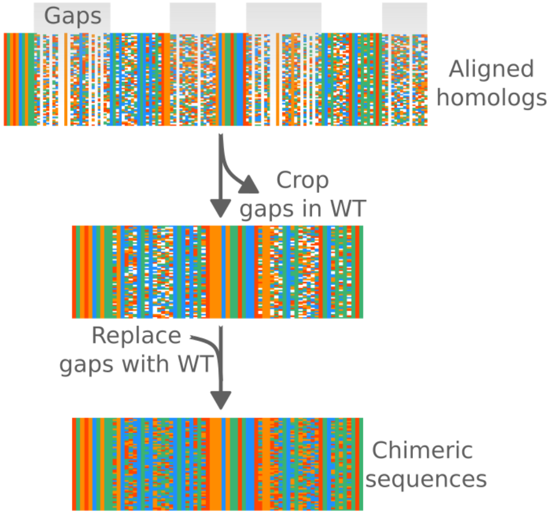


**Fig. S9. Design of the chimera sequences.** Homologs of Azo were found and aligned on the wild type sequence and secondary structure using the Infernal package. Insertions with respect to the wild type sequence were removed such that all sequences were all the same length (193 nucleotides). The deletions were then replaced by the wild type nucleotide at that position. We tested experimentally only the ones where at least 100 nucleotides were aligned to Azo (733 sequences).

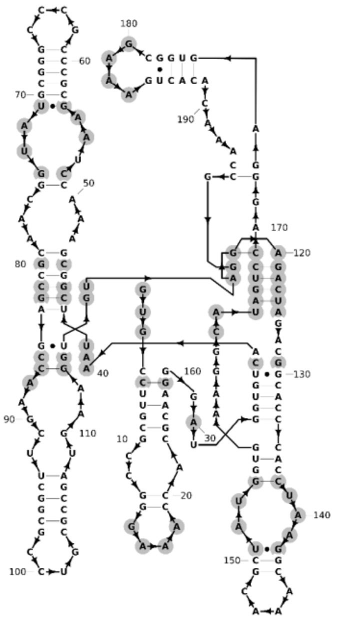


**Fig. S10. Tertiary constraints used with the BPR and SB models.** Gray positions correspond to positions that are in contact in the tertiary structure (*13*). These positions were held fixed in the wild type nucleotide when applying the 3D constraint design.

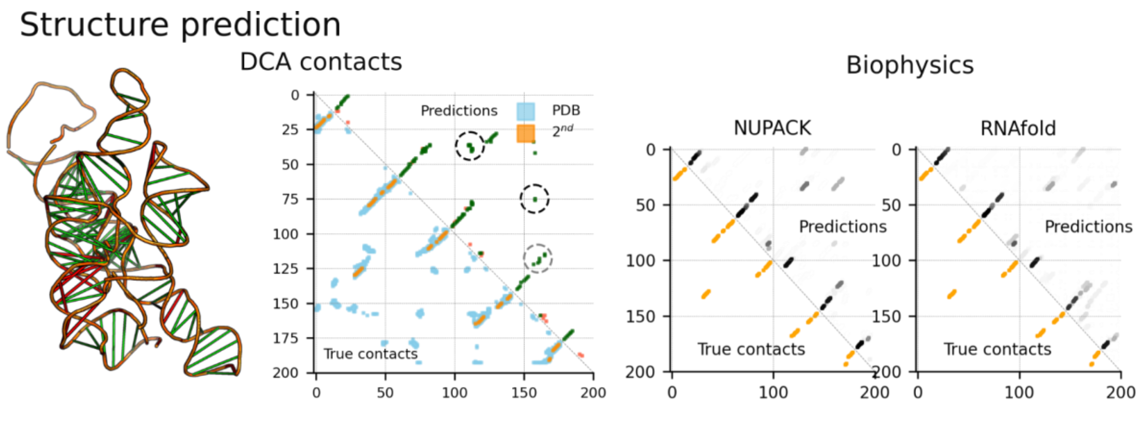


**Fig. S11. Structure prediction using evolutionary conservation and biophysical models.** On the left, we show the correct (green) and incorrect (red) predictions of contacts using DCA on the tertiary structure of Azoarcus X-ray structure 1G9B. The contact map inferred with DCA compared with the true tertiary and secondary contact also displays the predicted tertiary contacts (circled in gray). On the right, we show the base pair probability matrices predicted with NUPACK (which account explicitly for pseudoknot) and RNAfold (which do not account for pseudoknot).

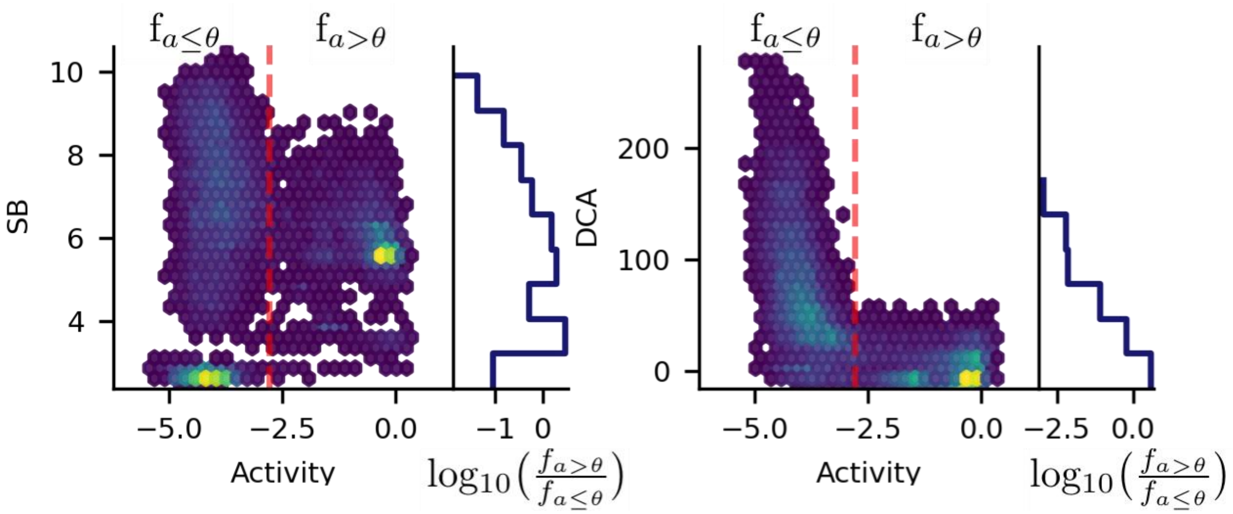


**Fig. S12. Structure-based (SB) score and evolutionary (DCA) score compared to the experimental activity.** Left: the graph shows the distribution of SB against the measured activity for the tested sequences where the color gradient represents the density of designs. The red dashed line represents the p-value=$10^{-3}$ threshold Θ=-2.76. The curve on the right displays the ratio between designs below and above the threshold . Right: Same but for the DCA score.

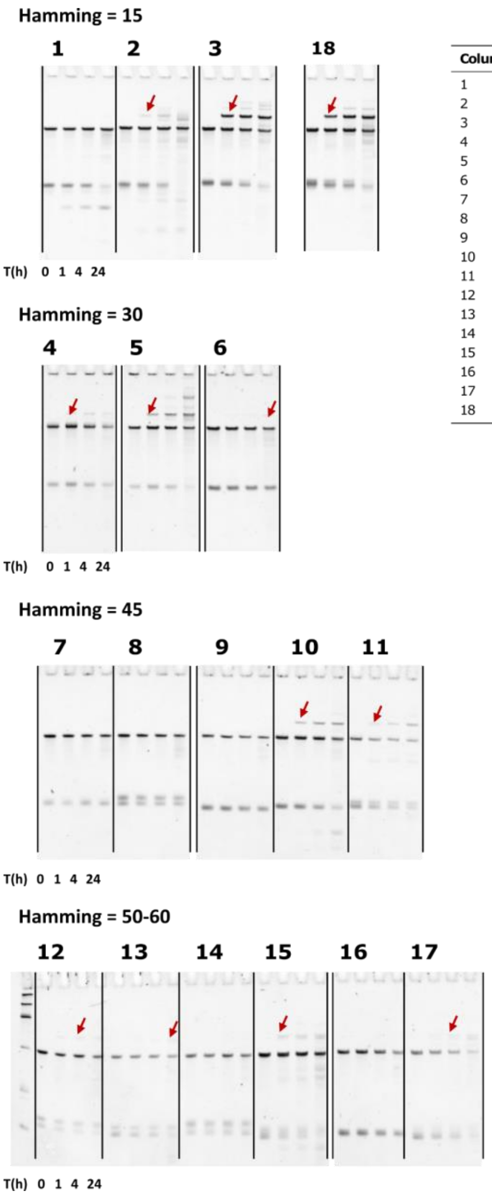


| Column | Sequence Name | Step 2 | Step 1+2 |
|---|---|---|---|
| 1 | dca.T1.15.2 | -0.431 | -2.494 |
| 2 | dca.T1.15.3 | -0.156 | -1.059 |
| 3 | dca.T1.15.4 | 0.511 | 0.479 |
| 4 | dca.T1.30.3 | -0.713 | -2.498 |
| 5 | dca.T1.30.5 | -0.375 | -0.397 |
| 6 | dca.T1.30.4 | 0.215 | -1.499 |
| 7 | dca.T1.45.3 | -0.788 | -2.168 |
| 8 | dca.T1.45.6 | -0.599 | -1.365 |
| 9 | dca.T1.45.4 | -0.075 | -1.168 |
| 10 | dca.T1.45.7 | -0.382 | -0.602 |
| 11 | dca.T1.45.5 | -0.321 | -0.357 |
| 12 | dca.T1.50.2 | -1.110 | -1.654 |
| 13 | dca.T1.50.1 | -0.612 | -1.293 |
| 14 | dca.T1.55.2 | -1.352 | -2.703 |
| 15 | dca.T1.55.1 | -0.358 | -1.522 |
| 16 | dca.T1.60.2 | 0.007 | -1.330 |
| 17 | dca.T1.60.1 | 0.570 | -1.232 |
| 18 | Azo | 0 | 0 |

**Fig. S13. Individual two-fragment self-reproduction assay for DCA designs. Self**-reproduction assay for 17 DCA designs. All designs tested were considered active (with an activity score above the activity threshold of -2.76). The table indicates the design name and their activity scores (step 2 refers to the ability of the design to take the substrate and step 1+2 refers to the ability of the design to remove the exon and then take the substrate). The red arrow shows the presence of the recombined covalent ribozyme.

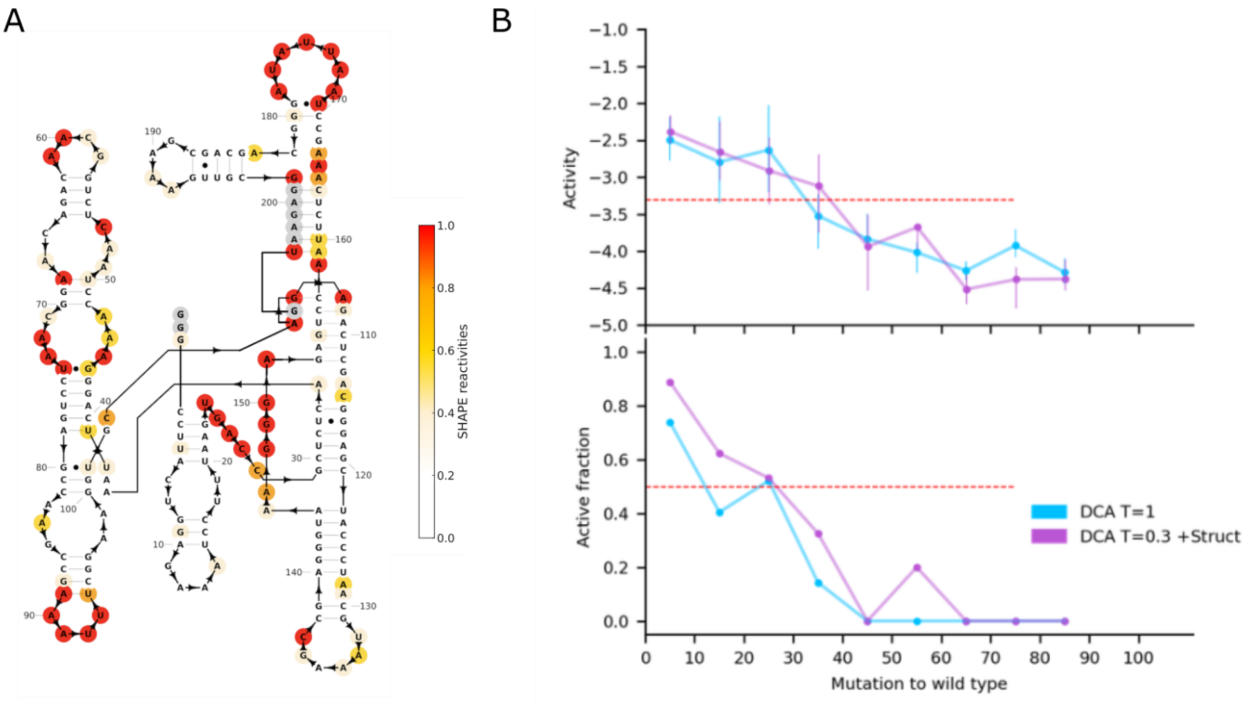


**Fig. S14. Self-splicing design and experimental activity measured for Phormidium variants.** (**A**) Shape reactivities projected on the 2D structure of Phordium group I intron, used for the structure-based generation, due to the lack of crystal structure. (**B**) The top panel shows the activity (relative to Azoarcus activity) across the number of mutations from the wild type Phormidium sequence. The dots represent the mean activity of each bin whereas the lines represent the first and third quartile of the activities. The dashed red line shows the activity threshold for p-value=$10^{-3}$. The bottom panel represents the active fraction— the fraction of designs above the activity threshold. The dashed line shows 0.5.

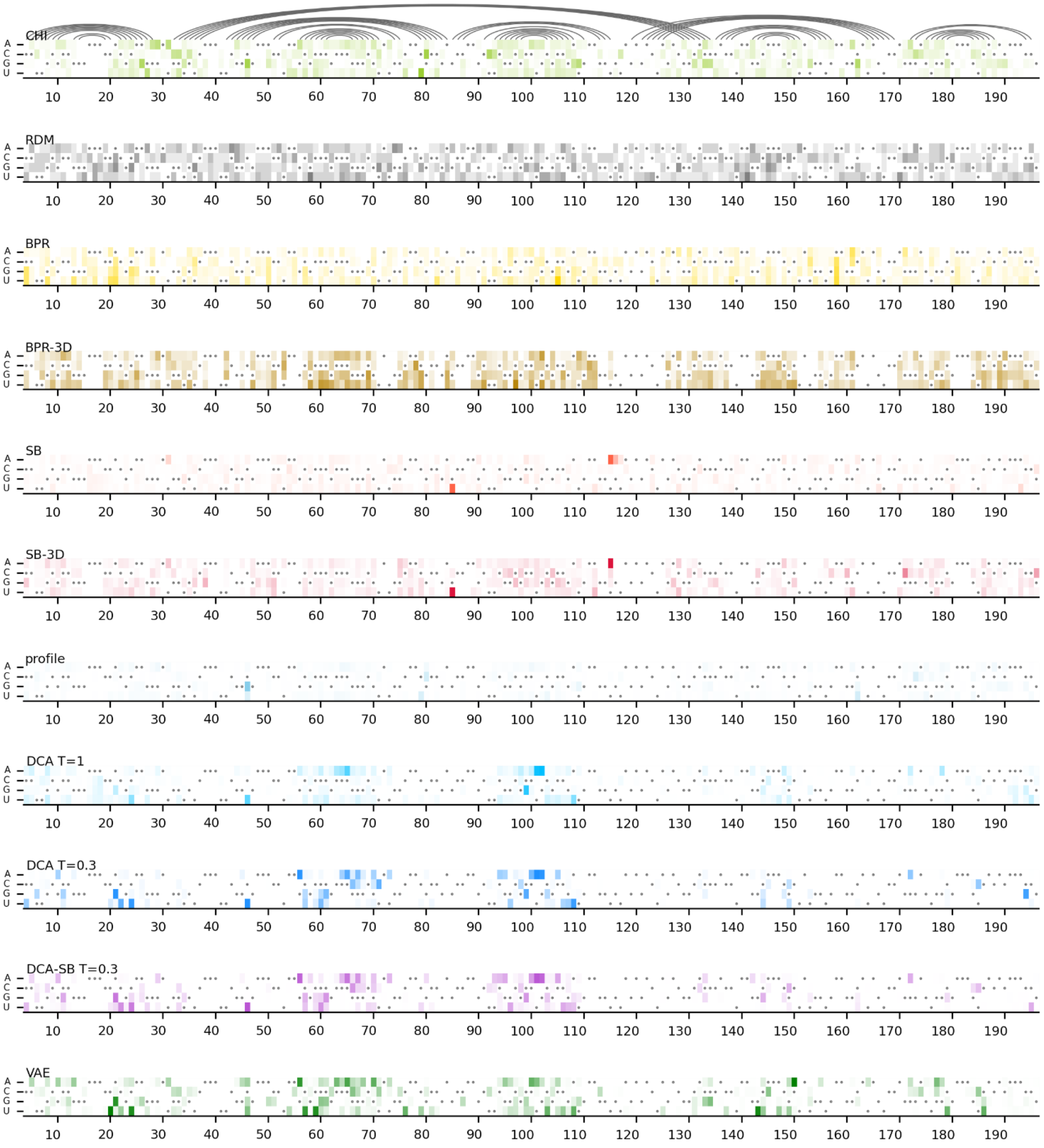


**Fig. S15. Distribution of mutations across the positions among active sequences per model.** Each row represents the frequency of mutations in all four nucleotides where dots represent the wild type nucleotides.

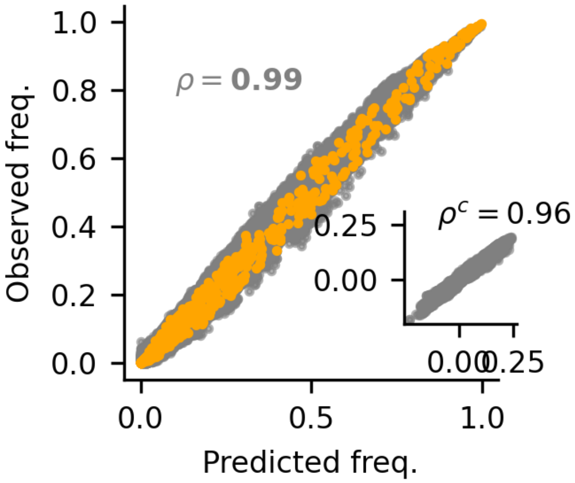


**Fig. S16. Reproduction of first- and second-order statistics with the DCA model.** The main scatter plot shows the correlation of single (orange) and pairwise (gray) frequencies between the MSA and the DCA predictions. The comparison with the prediction for the connected correlation $\rho_c = f_{ij} - f_i \cdot f_j$ is shown in the inserted scatter plot.

**Algorithm 1:** *bl-dca* implementation

```
Data: MSA, N, K Result: DCA
// Initialize the DCA parameters to zeros
DCA ← 0;
// Compute f_i and f_ij from the MSA
F ← frequencies(MSA);
for i ∈ 1 → N do
    // Pick a random sequence from the MSA
    s ← pick_one(MSA);
    // Perform K Monte Carlo steps to mutate the sequence s
    for k ∈ 1 → K do
        // Propose a mutation at a random position                he
        mut← mutate(s)
        // Apply the Metropolis criterion, and update s if
           accepted
        if Metropolis(mut) then s ← mut;
    // Update the DCA parameters
    for i ∈ 1 → L do
        // Update h_i
        for n_i ∈ {A, C, G, U,' −'} do
            DCA[i,n_i] -= (F[i,n_i] - δn_i, s_i) ×η;
        // Update J_ij
        for j ∈ i → L do
            for n_i ∈ {A, C, G, U,' −'} do
                for n_j ∈ {A, C, G, U,' −'} do
                    DCA[i,j,n_i,n_j] -= (F[i,j,n_i,n_j] - δn_i, s_i ×δn_j, s_j) ×η;
    // k is choosen such that η converges to 10^−4 in the end
    η ← η × k;
```

**Fig. S17. Pseudo-code of the BL-DCA implementation derived from (*7*).** The algorithm takes as argument the MSA, the number of parameter update steps N, and the number of MCMC steps. The learning parameter $\boldsymbol{\eta}$ is typically set to 0.001.

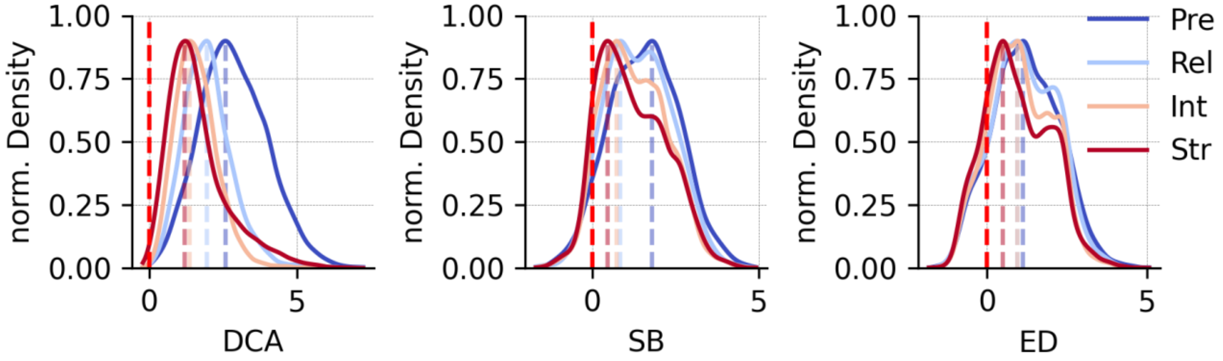


**Fig. S18. Validation of models on the benchmark mutational dataset.** Cross validation of the DCA, structure-based (SB), ensemble defect (ED) scores on an independent experimental dataset. Four experimental conditions were tested from the less stringent to the most (respectively in this order Pre, Rel, Int, and Str). The distributions represent the scores computed on the set of sequences obtained with that selection pressure, which we normalized such that the maximum (the mode) of the distribution is 0.9. The dashed lines show the position of the mode with respect to the wild type (defined at 0).

| name | # designs | # detected | # active designs | $L_{50}$ | fraction at $L_{50}$ | p at $L_{50}$ | $L_{max}$ | fraction at $L_{max}$ | p $L_{max}$ |
|---|---|---|---|---|---|---|---|---|---|
| **RDM** | 1687 | 1678 | 188 | 5 | 133/139 (95.7%) | < 1.00e-300 | 10 | 51/170 (30%) | 7.47e-110 |
| **BPR** | 1350 | 1344 | 156 | 15 | 41/69 (59.4%) | < 1.63e-104 | 20 | 4/76 (5.3%) | 1.21e-6 |
| **BPR+3D** | 1350 | 1348 | 231 | 15 | 32/59 (54.2%) | 4.72e-80 | 35 | 3/70 (4.3%) | 5.21e-5 |
| **SB** | 4200 | 4156 | 274 | 10 | 241/278 (86.7%) | < 1.00e-300 | 20 | 13/230 (5.7%) | 4.68e-19 |
| **SB-3D** | 1350 | 1347 | 231 | 10 | 129/147 (87.8)% | < 1.00e-300 | 40 | 3/99 (3.0%) | 1.46e-4 |
| **PRO** | 1800 | 1799 | 197 | 5 | 188/200 (94.0%) | < 1.00e-300 | 15 | 7/200 (3.5%) | 1.93e-9 |
| **DCA T=1** | 2547 | 2513 | 772 | 20 | 118/150 (78.7%) | < 1.00e-300 | 60 | 3/149 (2.0%) | 4.84e-4 |
| **DCA T=0.3** | 2203 | 2194 | 1344 | 45 | 66/104 (63.5%) | 3.48e-170 | 60 | 6/188 (3.2%) | 4.84e-8 |
| **DCA-SB** | 4200 | 4190 | 1634 | 55 | 150/175 (85.7%) | < 1.00e-300 | 65 | 8/116 (6.9%) | 5.77e-13 |
| **VAE** | 2800 | 2773 | 818 | 15 | 253/280 (90.4%) | < 1.00e-300 | 60 | 6/269 (2.2%) | 3.97e-7 |
| **CHI** | 733 | 729 | 46 | // | // | // | 65 | 5/96 (5.2%) | 5.67e-8 |
| **Total** | 24220 | 24071 | 5891 | | | | | | |

**Table S1 Models comparison.** Model acronym; Number of designs generated; Number of designs detected after sequencing; Number of designs with an activity score larger than the threshold; Highest mutation number of the $L_{50}$ bin ; Number of active designs over total number in the $L_{50}$ bin ; p-value for the 50% activity in the $L_{50}$ bin; Highest mutation number of the $L_{max}$ bin ; Number of active designs over total number in the $L_{max}$ bin ; p-value for the 50% fraction in the $L_{max}$ bin.

| distance → model ↓ | 5 | 10 | 15 | 20 | 25 | 30 | 35 | 40 | 45 | 50 | 55 | 60 | 65 | 70 | 75 | 80 |
|---|---|---|---|---|---|---|---|---|---|---|---|---|---|---|---|---|
| **RDM** | 11.71 | 20.96 | 29.08 | 36.42 | 43.17 | 49.42 | 55.25 | 60.71 | 65.82 | 70.62 | 75.13 | 79.37 | 83.35 | 87.07 | 90.55 | 93.80 |
| **PRO** | 9.48 | 16.90 | 23.39 | 29.14 | 34.46 | 39.28 | 43.70 | 47.63 | 51.56 | 55.01 | 58.25 | 61.20 | 63.96 | 66.44 | 68.65 | 70.82 |
| **RBP** | 10.17 | 17.91 | 24.55 | 30.45 | 35.79 | 40.68 | 45.20 | 49.38 | 53.27 | 56.90 | 60.30 | 63.47 | 66.44 | 69.20 | 71.78 | 74.18 |
| **RBP-3D** | 9.22 | 16.00 | 21.68 | 26.62 | 31.01 | 34.95 | 38.50 | 41.72 | 44.63 | 47.26 | 49.63 | 51.75 | 53.63 | 55.26 | 56.66 | 57.81 |
| **DCA T=1** | 7.04 | 12.58 | 17.28 | 21.41 | 24.94 | 28.11 | 30.83 | 33.37 | 35.57 | 37.52 | 39.29 | 40.62 | 41.87 | 43.03 | 43.96 | 44.36 |
| **DCA T=0.3** | 1.74 | 2.86 | 4.55 | 6.53 | 8.20 | 8.63 | 8.10 | | | | | | | | | |
| **DCA-SB** | 5.71 | 9.72 | 13.9 | | | | | | | | | | | | | |

**Table S2. Computed model support sizes. Values** are in $\log_{10}$ and are the theoretical values before correction by the experimental active fraction. Highlighted in bold is the support size corresponding to $L_{max}$. For the DCA-SB model, the support size is reported using the least-constrained parameter choice (alpha = 0.4). For the models DCA at T=0.3 and DCA-SB, support size estimation becomes infeasible in practice beyond a certain distance due to ergodicity issues. DCA T=1 support sizes have been reduced by a factor of ten because we only experimentally tested the sequences in the top 10% quantile of the DCA score.

| Sequence name | sequence | Distance from Azo |
|---|---|---|
| **Azo** | GUGCCUUGCGCCGGGAAACCACGCAAGGGAUGGUGUCAAA UUCGGCGAAACCUAAGCGCCCGCCCGGGCGUAUGGCAACGC CGAGCCAAGCUUCGGCGCCUGCGCCGAUGAAGGUGUAGAG ACUAGACGGCACCCACCUAAGGCAAACGCUAUGGUGAAGGC AUAGUCCAGGGAGUGGCGAAAGUCACACAAACCGG | 0 |
| **dca_sb_1434_60_70** | GUGCACUGCUCCGGGAAACCAAGUAGUGAAUGCUCUCAAA UUCAGGGAAACCUAAAUCUGGUAGUCCAGAUAAGGCAACC CUGAGCCAAGCCAAGUCACCUAUGACUUGGAAGGUGCAGA GACUCGACGGGAGCUACCUAACGGUUAGCCGAGGGUAAAG GGAGAGUCCAAUUACUGACGAAAGUCAGACAAAGAGG | 65 |
| **chimeric_466** | GUGGCUUGCGCCGGGAAACCACGCAAGCAAAUCAGGCUAA UUCGGGGAACGCCUAACGCCCGCCCGGGCGUAGGUCAAUCC CGAGCUAAGUCCCCGAUUUAAUUGGGGUAAAUGUGUAGAG ACUAUAUACCUGACACCAGUGGCAAACGCUAUGGUGAUGA GAUAGUCCAGUCCUUUUGGUAACAAGAGGAAACCGG | 65 |
| **chimeric_184** | GUGCCAUGCGCCGGGAAACCACGCAUGAUAACUUGGCUAA UUCGGGGAAGUCCAAGCGCCCGCCCGGGCGUAUGAUAAUCC CGAGCUAAAUUCUCCGCCAUGGCGGGGUAAAUGUGUAGAG ACUAUAUACCAGGGACCUAAAGCAAACGCCAAGGUCAUAA GAUAGUCCAAACCUUUAAGUAAUUAGAGGAAACCGG | 64 |
| **dca_f_mutations_213** | GUGCAUUCUGUUAGGAAACUUUAGAAUGAAUGGUGUCAAA UUCGGUGAAACCUAAGUCUUUGCCAAAAGAUAAGGCAACG CCGAGCCAAGCUCAUUUAGAAACAAUUGAGAAGGUGUAAC GACUAGACGGCACCCACCUAAGGUAAUCACAAUGGUGAAG GCAUAGUCUAGAGAGCAACCAAAGUUGCACAGGCACG | 60 |
| **dca_sb_1469_60_70** | GUGUAUGCCUGCGGGAAACCUAGGCAUGAAUGUGGUCAAA UUCGAUGAAACCUAAAUAGUGACAACACUAUAAGGCAAUA UCGAGCCAAGCCUUAUCAGUAAUGAUAAGGAAGGUGUAGA GACUAGACGGCCACCACCUAAAGGAAACCCUACGGUGAAGG UAUAGUCCAGAGAGUGGGGAAACUCACACGAACUGG | 60 |
| **dca_sb_1496_60_70** | GUGCAUUCAGGCAGGAAACUUCUGAGUGAAUGUGCUCAAA UUCGGUGAAACCUAAAGAGUGGAAACACUCUAAGGGAAUA CCGAGCCAAGCCUUUUCAUCAAUGAAAAGGAAGGUGUAGA GACUAGACGGGCACCACCUAAAGGAAAACCUAUGGUGAAG GUAUAGUCCAGAGAGUGGGGAAACUCACACAUACUGG | 60 |

| | | |
|---|---|---|
| **chimeric_382** | GUGCCUUCCGCCGGGAAACCACGGAAGAUUACUUGCCAAAUUCGGGGAAGCCCACGUUCCCGCCCGGGAACUAGGUAAUCCCGAGCUAAGCUCUGAUGUUUGUAUCAGAGAAAGUGUAGAGACUAGAUGGUAAGCACCUUAAAUAAUGAUUUAGGUGAAGGGAUAGUCCAGACUACAGCGAAAGUUGUAGAAACCGG | 62 |
| **dca_f_mutations_238** | GUGCUCUAAUAAGGGAAACCGAUUAGUGAAUGUGGUCAAAUUCAGGGAAGCUUAAGGAUAUUAUUAUAUCUAUGGUAACCCUGAGCCAAGCUUAGAAGCAAAUUUCUUUGAAGGUGCAGAGACUAGACGGCCACCACCUAAGGGAAACCCUAGGGUGAAGGGAUAGUCCAGGGAGUGACGAAAGUCACACGAAUUCG | 60 |

**Table S3. Active sequences at maximum distances**. Azo is the reference sequence. In this table, we only consider sequences that belong to bins where the active fraction is established with $p<0.001$. Sequences 'dca_sb_1434_60_70' and 'chimeric_466' are the ones found active at 65 mutations from Azo and are 99 mutations away from each other. The other pairs of sequences 99 mutations away from each other are: 'chimeric_466' and 'dca_f_mutations_213', 'chimeric_184' and 'dca_f_mutations_213', 'dca_sb_1469_60_70' and 'chimeric_184', 'dca_sb_1496_60_70' and 'chimeric_184'. The CHI 'chimeric_382' sequence is 59 mutations away from any other CHI sequence. The 'dca_f_mutations_238' generated by DCA at T=1 is 55 mutations away from any CHI sequence.

| Pool | [MgCl2] (mM) | Avg. length (nuc) | Nb. seq. |
|---|---|---|---|
| **Pre** | - | 173.9 | 12740 |
| **Rel** | 25 | 174.5 | 3346 |
| **Int** | 10 | 174.6 | 25586 |
| **Str** | 2 | 174.7 | 11803 |

**Table S4. Benchmark mutational dataset**. Size and sequence length of the four pools of sequences used for the DCA benchmarking with their average DCA score (relative to Azo), extracted from (*14*).

**Supplementary References:**

1. E. J. Hayden, N. Lehman, Self-assembly of a group I intron from inactive oligonucleotide fragments. *Chemistry & biology* **13**, 909–918 (2006).

2. P. L. Adams, M. R. Stahley, A. B. Kosek, J. Wang, S. A. Strobel, Crystal structure of a self-splicing group I intron with both exons. *Nature* **430**, 45–50 (2004).

3. I. Kalvari, E. P. Nawrocki, N. Ontiveros-Palacios, J. Argasinska, K. Lamkiewicz, M. Marz, S. Griffiths-Jones, C. Toffano-Nioche, D. Gautheret, Z. Weinberg, E. Rivas, S. R. Eddy, R. D. Finn, A. Bateman, A. I. Petrov. doi: https://doi.org/10.1093/nar/gkaa1047.

4. E. P. Nawrocki, S. Eddy, Infernal 1.1: 100-fold faster RNA homology searches. *Bioinformatics* **15;29(22):2933–5** (2013).

5. S. F. Altschul, W. Gish, W. Miller, E. W. Myers, D. J. Lipman, Basic local alignment search tool. *Journal of molecular biology* **215**, 403–410 (1990).

6. F. Calvanese, C. N. Lambert, P. Nghe, F. Zamponi, M. Weigt, Towards parsimonious generative modeling of RNA families. *Nucleic Acids Research* **52**, 5465–5477 (2024).

7. F. Cuturello, G. Tiana, G. Bussi, Assessing the accuracy of direct-coupling analysis for RNA contact prediction. *RNA* **26**, 637–647 (2020).

8. X. Ding, Z. Zou, C. L. Brooks III, Deciphering protein evolution and fitness landscapes with latent space models. *Nat Commun* **10**, 5644 (2019).

9. D. H. Mathews, M. D. Disney, J. L. Childs, S. J. Schroeder, M. Zuker, D. H. Turner, Incorporating chemical modification constraints into a dynamic programming algorithm for prediction of RNA secondary structure. *Proceedings of the National Academy of Sciences* **101**, 7287–7292 (2004).

10. M. Zuker, P. Stiegler, Optimal computer folding of large RNA sequences using thermodynamics and auxiliary information. *Nucleic Acids Res* **9**, 133–148 (1981).

11. J. S. McCaskill, The equilibrium partition function and base pair binding probabilities for RNA secondary structure. *Biopolymers: Original Research on Biomolecules* **29**, 1105–1119 (1990).

12. R. M. Dirks, N. A. Pierce, An algorithm for computing nucleic acid base-pairing probabilities including pseudoknots. *J Comput Chem* **25**, 1295–1304 (2004).

13. A. M. Mustoe, H. M. Al-Hashimi, C. L. Brooks, Secondary structure encodes a cooperative tertiary folding funnel in the Azoarcus ribozyme. *Nucleic Acids Research* **44**, 402–412 (2016).

14. E. J. Hayden, D. P. Bendixsen, A. Wagner, Intramolecular phenotypic capacitance in a modular RNA molecule. *Proceedings of the National Academy of Sciences* **112**, 12444–12449 (2015).

15. T. M. Cover, J. A. Thomas, *Elements of Information Theory* (Wiley, New York, 1999).
